# DGKα and ζ Deficiency Causes Regulatory T-Cell Dysregulation, Destabilization, and Conversion to Pathogenic T-Follicular Helper Cells to Trigger IgG1-Predominant Autoimmunity

**DOI:** 10.1101/2024.11.26.625360

**Authors:** Lei Li, Hongxiang Huang, Hongxia Wang, Yun Pan, Huishan Tao, Shimeng Zhang, Peer W.F. Karmaus, Michael B. Fessler, John W. Sleasman, Xiao-Ping Zhong

## Abstract

Regulatory T cells (Tregs) actively engage in immune suppression to prevent autoimmune diseases but also inhibit anti-tumor immunity. Although Tregs express a TCR repertoire with relatively high affinities to self, they are normally quite stable and their inflammatory programs are intrinsically suppressed. We report here that diacylglycerol (DAG) kinases (DGK) ( and ( are crucial for homeostasis, suppression of proinflammatory programs, and stability of Tregs and for enforcing their dependence on CD28 costimulatory signal. Treg-specific deficiency of both DGK( and ( derails signaling, metabolic, and transcriptional programs in Tregs to cause dysregulated phenotypic and functional properties and to unleash conversion to pathogenic exTregs, especially exTreg-T follicular helper (Tfh) 2 cells, leading to uncontrolled effector T cell differentiation, deregulated germinal center (GC) B-cell responses and IgG1/IgE predominant antibodies/autoantibodies, and multiorgan autoimmune diseases. Our data not only illustrate the crucial roles of DGKs in Tregs to maintain self-tolerance but also unveil a Treg-to-self-reactive-pathogenic-exTreg-Tfh-cell program that is suppressed by DGKs and that could exert broad pathogenic roles in autoimmune diseases if unchecked.

## Introduction

Regulatory T cells (Tregs) are crucial for maintaining self-tolerance to prevent autoimmune diseases (1–3). The Treg-specific transcription factor (TF) Foxp3 is crucial for Treg generation, identity, and function (4, 5). Natural Tregs are generated in the thymus, express TCRs with relatively high affinities to self-antigens, and display a certain degree of self-reactivity. However, Tregs express suppressive cytokines but not IL-2 and other proinflammatory cytokines associated with T helper cells (6). Foxp3 and Treg-specific developmentally established epigenetic landscapes suppress overt proinflammatory responses in Tregs (7–9). Tregs can be pathogenic for autoimmune diseases if they lose suppressive functions, gain proinflammatory functions, and/or become unstable and convert to exTregs because of their self-reactivity (10–13). However, mechanisms that control Treg stability and function are still not fully understood.

Autoantibodies are hallmarks for many autoimmune diseases. GC is a major site of Ig class-switch and antibody affinity maturation and as such are essential for protective humoral immune response and, unfortunately, development of autoantibodies (14, 15). Tfh cells promote GC B-cell proliferation and survival, Ig-class switch and affinity maturation, and memory B-cell and long-lived plasma cell formation (15). However, deregulated Tfh-cells can trigger abnormal GC B and memory B-cell responses, leading to augmented autoantibody production (16, 17). T follicular regulatory (Tfr) cells, a specialized Treg sublineage expressing CXCR5, PD-1, and the TF Bcl6, suppress GC responses including autoantibody responses (18, 19). Abnormal Tfh and Tfr-cells have been associated with or are the causal factors of autoimmune diseases in humans and animals (17, 19, 20). However, the origins of pathogenic self-reactive Tfh-cells in autoimmune diseases are ellusive.

TCR signal participates in Foxp3 induction, Treg-specific CpG hypomethylation formation, and Treg maintenance (21, 22). A critical event after TCR engagement is PLCγ1-mediated generation of two important second messengers, DAG and inositol tris-phosphates. DAG associates with and activates multiple effector molecules such as RasGRP1 and PKCθ, leading to activation of the Ras-Erk1/2 and IKKα/β/γ-NFκB pathways as well as PI3K/Akt-mTOR signaling (23). DAG can be controlled by DGKs, a family of ten isoforms phosphorylating DAG to produce phosphatidic acid (24, 25). DGKα and DGKζ, the major isoforms expressed in T-cells, regulate DAG-mediated RasGRP1-Ras-Erk1/2, PKCθ-IKK-NFκB, and PI3K/Akt-mTOR pathways in T cells to control their development, activation, anergy, survival, effector function, and antimicrobial and antitumor immunity as well as *i*NKT and MAIT-cell development (23, 26–37). Although deficiency of DGKζ alone facilitated Treg development (38, 39), whether DGKα and ζ may function synergistically in Tregs, and their true in vivo importance in Tregs for self-tolerance, have remained unclear.

We report here that DGKα and ζ synergistically ensure proper signaling, metabolic, and transcriptional programs in Tregs to ensure their normal homeostasis, stability, and lack of inflammatory property and to enforce their dependence of CD28 costimulatory signal. DGKα and ζ double-deficiencies (DKO) in Tregs unleashes their proinflammatory programs, alleviates CD28-dependence for their development and homeostasis, and facilitates their conversion to pathogenic exTregs, especially exTreg-Tfh2-cells, causing multiorgan autoimmune diseases, deregulated GC B-cell responses and IgG1/IgE predominant autoantibodies, and lupus-like diseases.

## RESULTS

### Treg-specific DGKα and ζ double deficiency causes multiorgan autoimmune diseases

To determine the role of DGKα and ζ in Tregs for self-tolerance, we analyzed *Dgka^-/-^z^f/f^-Foxp3^YFPCre/YFPCre^*Treg-specific DGKαζDKO (DKO-Cre or Treg-αζDKO) and control WT*-Foxp3^YFPCre/YFPCre^* (WT-Cre) mice. Treg-αζDKO mice had weight losses (Figure 1A) and developed lymphoproliferative disorders and multiorgan autoimmune diseases, manifested by enlarged spleen and lymph nodes (LNs) with increased total cellularity (Figure 1B,1C), mononuclear cell infiltration in the liver, lung, kidney, thyroid gland, and pancreas, significant thickening of the epidermis of skin (Figure 1D, Supplemental Figure S1), and elevated anti-double strand DNA (dsDNA), -single strand DNA (ssDNA), and -nuclear autoantibodies (Figure 1E, 1F). Elevated autoantibodies were mainly caused by increased IgG1 but not IgG2b or IgG3 (Figure 1G), suggesting enhanced type-2 autoimmunity. In the kidney, thickened glomerular basal membrane (arrows in Figure 1D) and IgG deposition in the glomeruli could be observed (Figure 1H). Although DGKα and ζ can function individually to promote T cell anergy and enhance T cell activation in certain experimental settings, neither DGKζ nor DGKα deficiency caused obvious lymphoproliferative or autoimmune diseases (Supplemental Figure S2). Thus DGKα and ζ function synergistically in Tregs to ensure self-tolerance.

**Figure 1.**
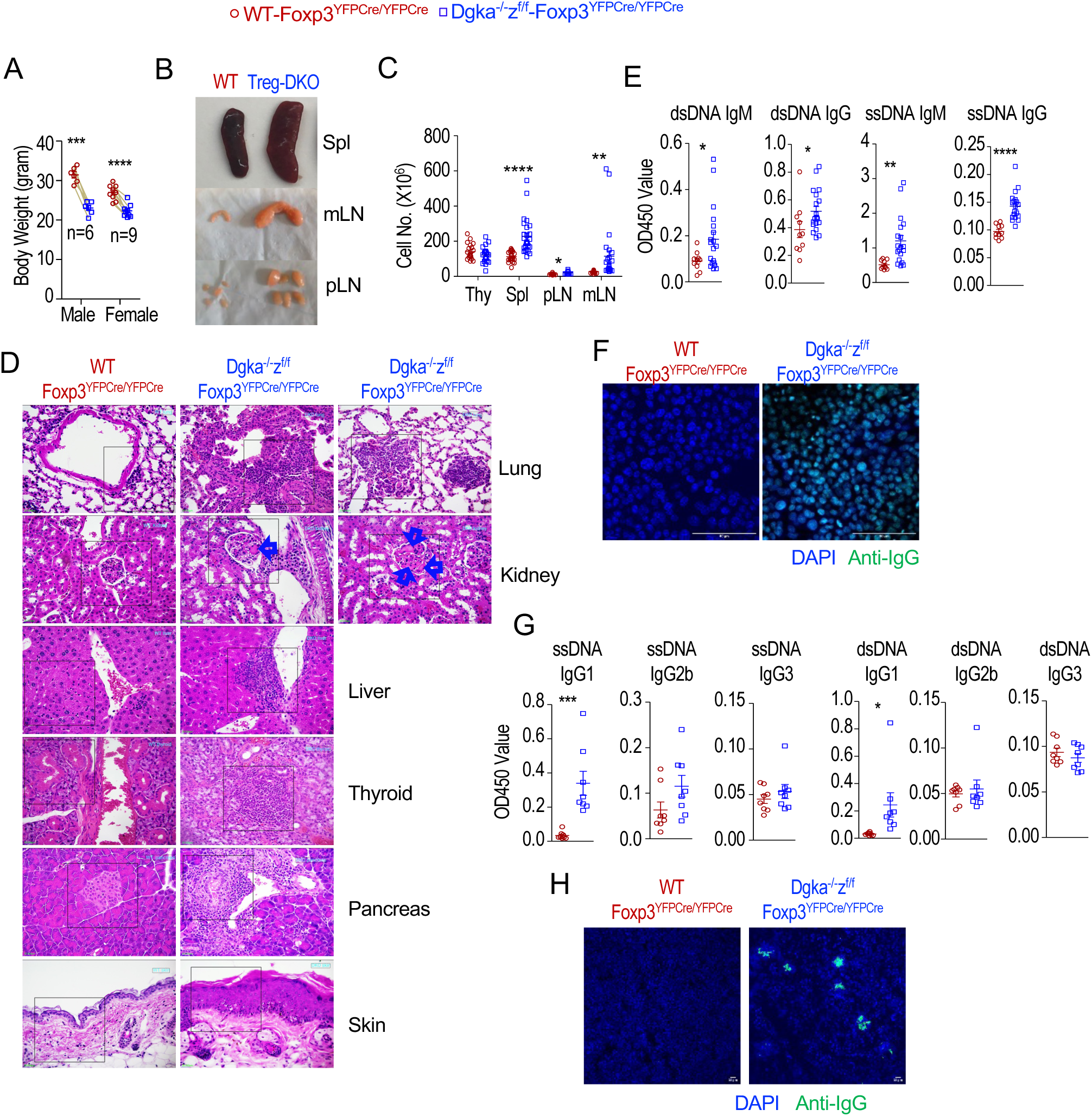
Autoimmune diseases in Treg-αζDKO mice. Female *Dgka^-/-^z^f/f^-Foxp3^YFPCre/YFPCre^*or male *Dgka^-/-^z^f/f^-Foxp3^YFPCre^* and WT control mice were analyzed. **A.** Body weights of 5–9-month-old mice. **B.** Representative pictures of indicated organs in a pair of 6-month-old mice. **C.** Total cell numbers in the indicated organs in 2–9-month-old mice. **D.** Representative H&E staining of paraffin thin-sections of indicated organs. **E.** Seral autoantibodies titers in 7-month-old mice. **F.** Representative images of detection of seral antinuclear antibodies against fixed HEp-2 cells. **G.** Seral autoantibody IgG subtypes. **H.** Detection of IgG deposition in cyro-sections of kidneys with fluorescence confocal microscopy. Data shown are representative of or pooled from at least six experiments. Each circle or square represents one mouse of the indicated genotypes. Each line connecting the circle and square represents one pair of age- and sex-matched mice examined in one experiment. *, p < 0.05; **, p < 0.01; ***, p < 0.001; ****, p < 0.0001 determined by two-tail pair-wise (Figure 1A, data with lines connecting WT and αζDKO mice) or unpaired Student *t* test.

### Treg-specific DGK**αζ**DKO enhances Treg homeostatic expansion

In Treg-αζDKO mice, CD4^+^Foxp3^+^ Tregs and CD4^+^CD25^+^Foxp3^-^ pre-Tregs were not altered in the thymus (Figure 2A, 2B) but increased in the spleen and LNs (Figure 2A, 2C). αζDKO Tregs survived similarly (Figure 2D) but increased in proliferation as indicated by Ki67 expression (Figure 2E) and BrdU incorporation (Figure 2F). They expressed slightly increased TF Helios, suggesting they were tTregs, although Nrp1 expression was similar or slightly reduced (Figure 2G). In contrast, Treg percentages and numbers were not altered in *Dgka^-/-^-Foxp3^YFPCre/YFPCre^*and *Dgka^+/+^z^f/f^-Foxp3^YFPCre/YFPCre^* mice (Supplemental Figure S3A–S3D).

**Figure 2.**
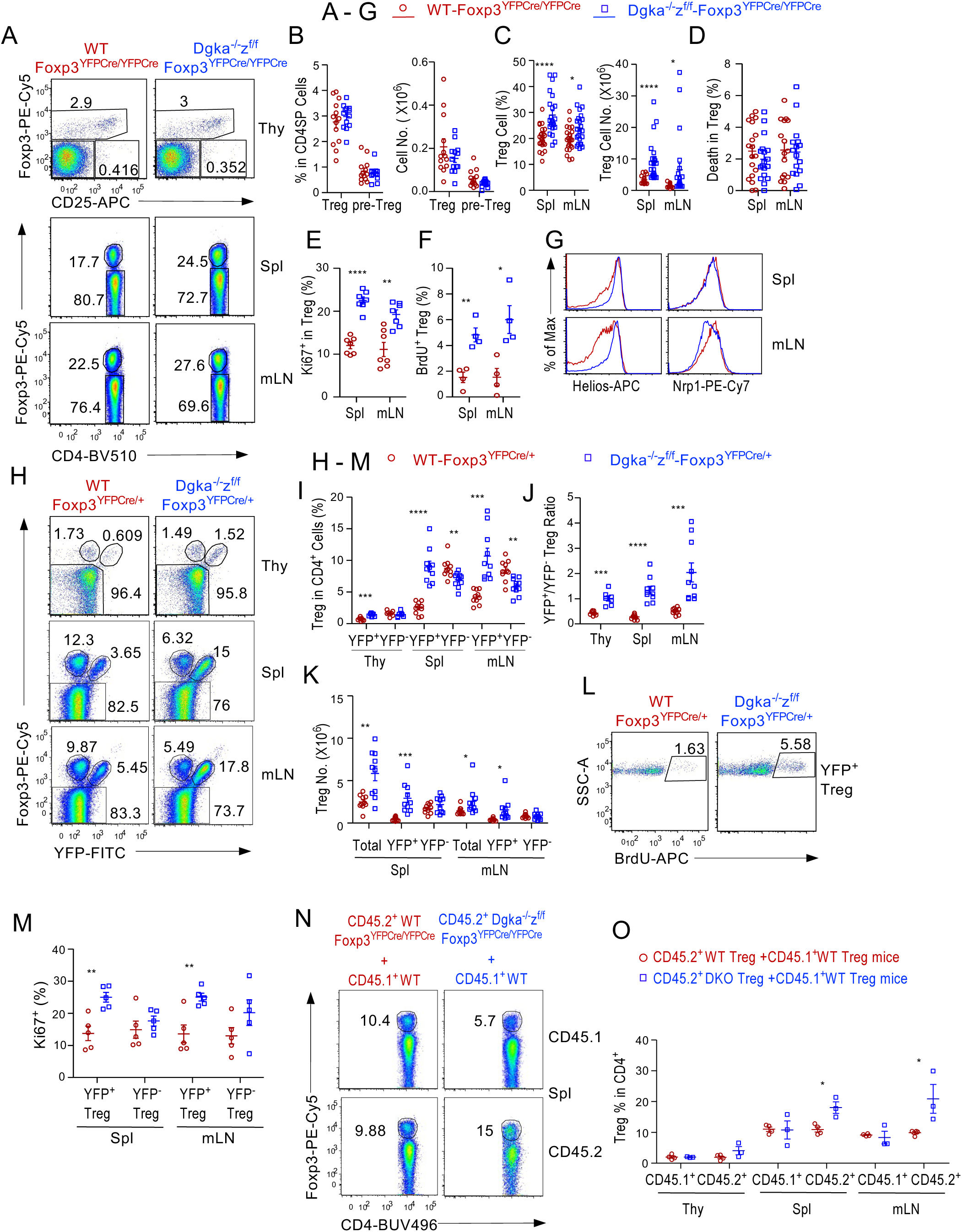
Enhanced Treg expansion in Treg-αζDKO mice. **A–G.** Analysis of *Dgka^-/-^z^f/f^-Foxp3^YFPCre/YFPCre^*and WT*-Foxp3^YFPCre/YFPCre^* control mice. **A.** Representative FACS plots showing Foxp3 and CD25 staining in CD4^+^ SP thymocytes (top panels) and CD4 and Foxp3 staining in splenic and LN CD4^+^ T cells. **B.** Foxp3^+^ Treg and CD25^+^Foxp3^-^CD4^+^ pre-Treg percentages and numbers in the thymus. **C.** Foxp3^+^ Treg percentages and numbers in the spleen and mLNs. **D.** Death rate of Tregs. **E.** Percentages of Ki67^+^ cells within Tregs. **F.** BrdU incorporation in Tregs 8–10 hours after intraperitoneal (*i.p.*) injection of BrdU. **G.** Overlaid histograms showing Helios and Nrp1 expression in Tregs. **H–M.** Analysis of female *Dgka^-/-^z^f/f^-Foxp3^YFPCre/+^*and WT-*Foxp3^YFPCre/+^* mice. **H.** Representative FACS plots showing intracellular Foxp3 and YFP staining in CD4^+^ T cells. **I.** YFP^+^Foxp3^+^ and YFP^-^Foxp3^+^ Treg percentages in CD4^+^ T cells. **J.** YFP^+^Foxp3^+^/YFP^-^Foxp3^+^ ratios in individual mice. **K.** Total, YFP^+^, and YFP^-^ Treg numbers in the spleen and mLNs. **L.** Representative FACS plots showing BrdU incorporation in LN YFP^+^ Tregs. **M.** Percentages of Ki67^+^ cells in YFP^+^ and YFP^-^ Tregs. **N, O.** Analyses of mixed BM chimeric mice. CD45.1^+^CD45.2^+^ WT mice were lethally irradiated and intravenously (*i.v.)* injected with a mixture of CD45.1^+^ WT with either CD45.2^+^ WT-*Foxp3^YFPCre/YFPCre^*or *Dgka^-/-^z^f/f^-Foxp3^YFPCre/YFPCre^* BM cells. Recipient mice were analyzed 6–8 weeks after reconstitution. **N.** Intracellular staining of Foxp3 in CD4^+^TCRβ^+^ T cells. **O.** CD45.1^+^CD45.2^-^ and CD45.1^-^CD45.2^+^ Treg percentages in individual mice. Data shown are representative of or pooled from 5–23 experiments except F, L, N, and O. F is pooled from four experiments. L represents two experiments. N and O are representative or pooled from three experiments. *, p < 0.05; **, p < 0.01; ***, p < 0.001; ****, p < 0.0001 determined by two-tail unpaired Student *t* test.

In female *Dgka^-/-^z^f/f^-Foxp3^YFPCre/+^*(DKO-Cre^het^) mice that contained both YFP^-^ Foxp3^+^*Dgka^-/-^z^wt^* Tregs to limit autoimmunity and YFP^+^Foxp3^+^αζDKO Tregs, YFP^+^Foxp3^+^ Treg percentages were increased in the thymus, spleen, and LNs, while YFP^-^Foxp3^+^ Tregs were decreased in the spleen and LNs (Figure 2H, 2I), leading to increased YFP^+^/YFP^-^ Treg ratios (Figure 2J). Moreover, female αζDKO-Cre^het^ mice had increased total and YFP^+^ but not YFP^-^ Treg numbers (Figure 2K) owing to hyperproliferation of YFP^+^ but not YFP^-^ Tregs (Figure2L, 2M, Supplemental Figure S3E). Additionally, YFP^+^αζDKO Tregs also expressed similar or increased levels of tTreg marker Helios (Supplemental Figure S3F).

In lethally irradiated CD45.1^+^CD45.2^+^ recipient mice reconstituted with a mixture of CD45.1^+^ WT BM cells with either CD45.2^+^ WT or *Dgk*a*^-/-^z^f/f^-Foxp3^YFPCre/YFPCre^*BM cells, CD45.2^+^ Treg-αζDKO-derived Treg percentages within CD4^+^TCRβ^+^ cells were increased compared with CD45.1^+^ WT-derived Treg cells in the same recipient mice as well as with CD45.2^+^ WT-derived Tregs in CD45.2^+^ WT/CD45.1^+^ WT BM chimeric mice (Figure 2N, 2O).

Together, these data demonstrate that DGKα and ζ play an important, synergistic, and intrinsic role for Treg homeostasis by inhibiting Treg proliferation.

### Altered signaling, metabolic, and mTOR pathways and Th signatures in Tregs in Treg-αζDKO mice

To understand how DGKαζDKO deregulated Tregs, we first examined signaling and metabolism in αζDKO-Tregs. αζDKO CD62L^+^CD44^low^ central (c) and CD62L^-^CD44^+^ effector (e) Tregs displayed increased Erk1/2, S6, and AktS473 phosphorylation (indications of enhanced Ras-Erk1/2, mTORC1, and mTORC2 signaling, Figure 3A); elevated nutrient transporters CD71 and CD98 (Figure 3B); increased glucose uptake (Figure 3C); and enhanced glycolysis (Figure 3D, 3E). Thus, DGKα and ζ inhibited DAG-mediated Ras-Erk1/2 and mTOR signaling in Tregs and were important for proper Treg metabolism.

**Figure 3.**
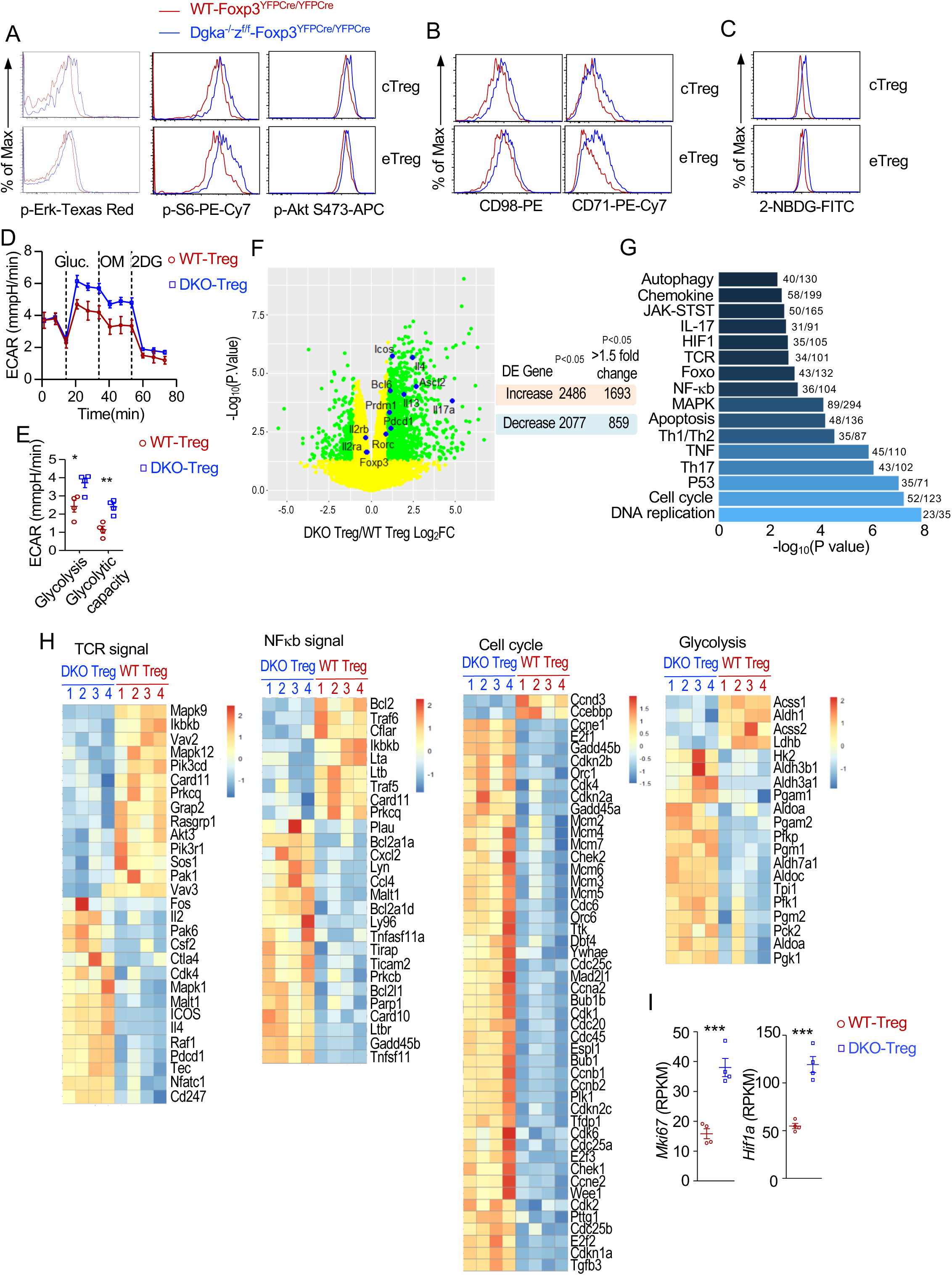
Altered signaling, metabolism, and transcriptional programs in αζDKO Tregs. Tregs in peripheral lymphoid organs from *Dgka^-/-^z^f/f^-Foxp3^YFPCre/YFPCre^*and WT*-Foxp3^YFPCre/YFPCre^* control mice were analyzed. **A, B.** Overlaid histograms show Erk1/2, S6, and Akt S473 phosphorylation (A) and CD71 and CD98 expression (B) in cTregs and eTregs. **C.** Overlaid histograms show 2-NBDG uptake in Tregs. **D, E.** Extracellular acidification rate (ECAR) measurements of sorted Tregs (n = 4 for both WT and DKO Tregs) following sequential treatment with glucose, oligomycin (OM, for mitochondrial perturbation), and 2DG (a glucose inhibitor). Representative ECAR profiles (D) and summary scatter plot of ECAR (E). **F.** Volcano plot comparison of gene expression between WT and DKO Tregs. Green-colored genes are differentially expressed with greater than 1.5-fold differences between WT and αζDKO Tregs (p < 0.05). Right panel shows total numbers of DEGs in DKO Tregs. **G.** Prominently changed KEGG pathways. **H.** Heatmaps show DEGs in TCR signaling, NFκB, cell cycle, and glycolysis. **I.** *Hif1a* and *Mki67* mRNA levels. *, p < 0.05; **, p < 0.01; ***, p < 0.001 determined by two-tail unpaired Student *t* test.

We next performed transcriptomic analyses after sequencing RNA from WT*-* and *Dgka^-/-^ z^f/f^-Foxp3^YFPCre/YFPCre^*CD4^+^Foxp3YFP^+^ Tregs. There were 4,563 differentially expressed genes (DEGs, p<0.05) between these Tregs, with 2,486 upregulated (1,693 > 1.5-fold increase) and 2,077 downregulated (859 >1.5-fold decrease) in αζDKO-Tregs (Figure 3F, Supplemental Table 1). KEGG pathway analyses revealed 132 enriched pathways (Figure 3G, Supplemental Table 2), including those involved in cell expansion and homeostasis (DNA replication, cell cycle, P53, apoptosis, and autophagy pathways), Th differentiation and function (Th1/2/17, IL17, Foxo, cytokines and cytokine signaling pathways, and Tfh pathway to be shown in Supplemental Figure S7), TCR signaling and T cell activation (TCR, NFκB, MAPK, and cytokine pathways), metabolism (glycolysis, pyruvate, citrate cycle, glutathione, purine metabolism, fatty acid pathways, and nutrient transporters), and cell migration and homing (chemokines and their receptors, Figure 3H, Supplemental Figure S4). αζDKO-Tregs upregulated many cell cycle-related genes including *Mki67* (Figure 3I), indicating that DGKαζ inhibited the cell cycle machinery to limit the normal Treg pool size. The enrichment of the glycolysis and Hif1 pathways, including upregulation of rate-limiting enzymes *Hk2* and *Pfk1* and *Hif1a* in DKO-Tregs (Figure 3G–3I), suggested that DGKαζ inhibited glycolysis via multiple mechanisms.

In αζDKO-Tregs, several critical TCR signaling components such as *Vav2/3*, *Rasgrp1*, *Prkcq*, *Ikbkb, Card11*, *Pik3cd*, *Pik3r1*, *Akt3, and Pak1*, were decreased, likely as a result of negative feedback mechanisms. Several other signal components, such as *Cd247* (encoding CD3ζ), *Tec*, *Raf1*, *Mapk1*, and *Malt1*, were increased. The enrichment of both MAPK and NFκB pathways (Figure 3H, Supplemental Figure S4), coupled with enhanced Erk1/2 phosphorylation and mTORC1/2 activation (Figure 3A), indicated that DGKα and ζ prevented dysregulated DAG-mediated signaling in Tregs.

### Dysregulated eTreg differentiation, enhanced response to TCR stimulation, and altered properties of Treg-**αζ**DKO Tregs

Differentiation of cTregs to eTregs is important for immune suppression. In *Dgka^-/-^z^f/f^-Foxp3^YFPCre/YFPCre^*mice, cTreg and eTreg percentages were decreased and increased, respectively (Figure 4A, Supplemental Figure S5A). Similar changes were also observed in YFP^+^ but not YFP^-^ Tregs in female *Dgka^-/-^z^f/f^-Foxp3^YFPCre/+^* mice (Figure 4B, Supplemental Figure S5B) and in CD45.2^+^ *Dgka^-/-^z^f/f^-Foxp3^YFPCre/YFPCre^*Tregs in mixed BM chimeric mice (Figure 4C, 4D). The increased eTreg but decreased cTreg percentages were not due to decreases of cTreg numbers as cTreg numbers were not decreased in *Dgka^-/-^z^f/f^-Foxp3^YFPCre/YFPCre^*mice or even increased within YFP^+^ splenic Tregs in female *Dgka^-/-^z^f/f^-Foxp3^YFPCre/+^*mice (Figure 4A, 4B). Moreover, both cTregs and eTregs from *Dgka^-/-^z^f/f^-Foxp3^YFPCre/YFPCre^*mice showed enhanced proliferation (Supplemental Figure S5C, S5D). The disproportional increases of αζDKO eTregs might be caused by accelerated cTreg to eTreg differentiation in the absence of both DGKα and ζ. However, a contribution of increased proliferation of αζDKO eTreg proliferation could not be ruled out. In contrast to Treg-αζDKO mice, c/eTreg ratios were not obviously altered in *Dgka^-/-^*-or *Dgka^+/+^z^f/f^-Foxp3^YFPCre/YFPCre^*mice (Supplemental Figure S6A–S6D). Thus, DGKα and ζ intrinsically and synergistically inhibited cTreg to eTreg differentiation.

**Figure 4.**
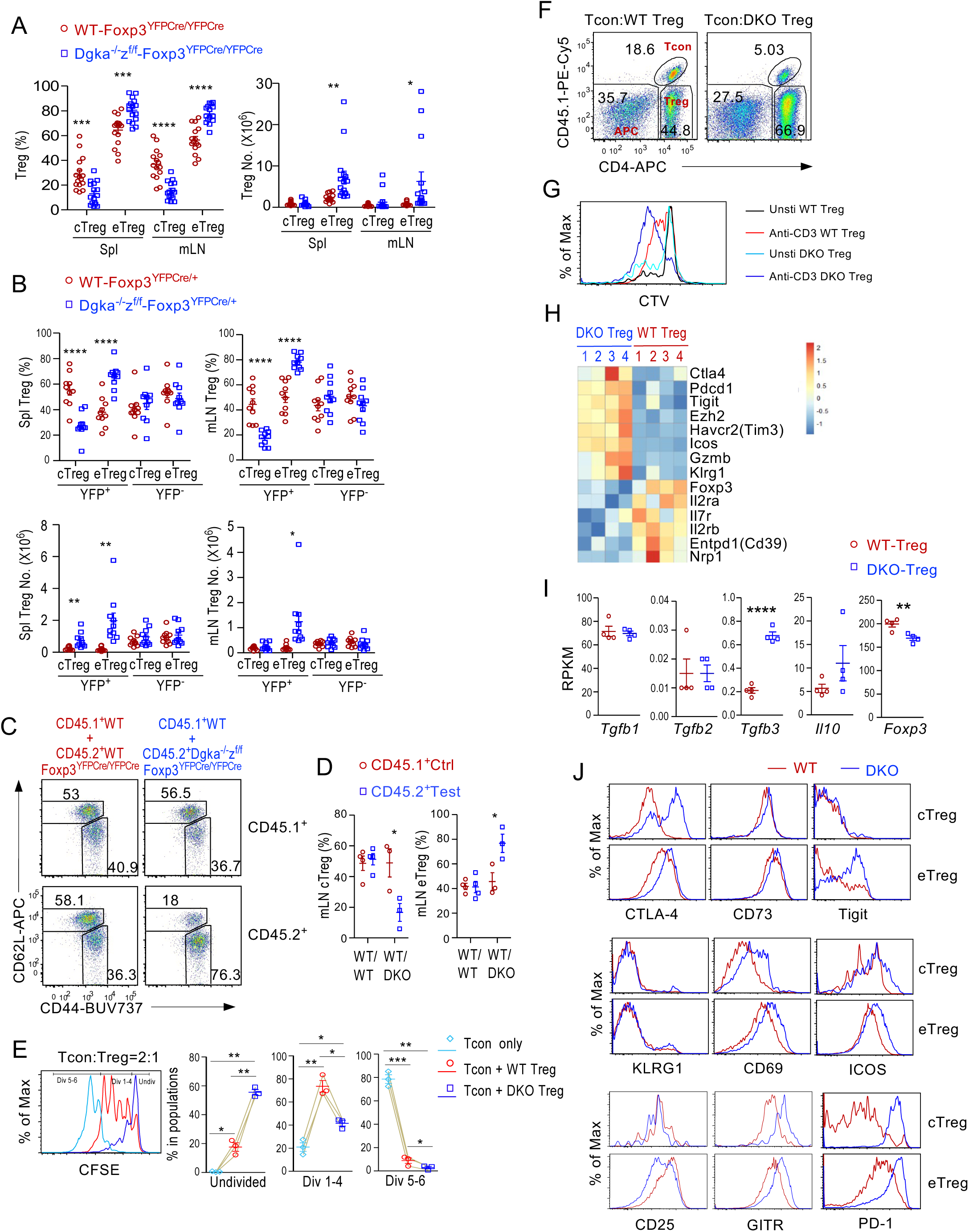
Enhanced effector differentiation and altered properties of αζDKO Tregs. **A.** Scatter plots show mean ± SEM of cTreg and eTreg percentages and numbers in *Dgka^-/-^z^f/f^-Foxp3^YFPCre/YFPCre^* and WT-*Foxp3^YFPCre/YFPCre^* mice. **B.** Scatter plots show mean ± SEM of cTreg and eTreg percentages and numbers of YFP^+^ and YFP^-^ CD4^+^Foxp3^+^ Tregs in female *Dgka^-/-^z^f/f^-Foxp3^YFPCre/+^* and WT*-Foxp3^YFPCre/+^* mice. **C, D.** Analyses of mixed BM chimeric mice as described in Figure 2N. **C.** CD44 and CD62L expression in CD45.1^+^ WT and in CD45.2^+^ WT or αζDKO Tregs in mLNs. **D.** Scatter plots show mean ± SEM of cTreg and eTreg percentages in CD45.2^+^ and CD45.1^+^ Tregs. **E, F.** In vitro contact inhibition assay. CTV labeled WT CD45.1^+^CD4^+^Foxp3YFP^-^ Tcon were mixed with 2:1 ratio of CD45.1^-^CD45.2^+^ WT or αζDKO Tregs in the presence of mitomycin C treated splenocytes as antigen presenting cells (APCs) from TCRα^-/-^ mice and were stimulated with an anti-CD3 antibody for 72 hours. **E.** Overlaid histograms show CTV dilution of CD45.1^+^ Tcon. Scatter plots show mean ± SEM of Tcon cells that were undivided, divided 1 – 4 times, and divided >5 times. **F.** Tcon, Treg, and APC populations revealed by CD45.1 and CD4 expression. **G.** TCR-induced Treg proliferation in vitro. Overlaid histogram showing CTV dilution of Foxp3YFP^+^ Tregs from CTV-labeled splenocytes after anti-CD3 stimulation for 72 hours. **H.** Heatmap showing differentially expressed Treg effector molecules between WT and αζDKO Tregs (p < 0.05). **I.** mRNA levels of *Il10*, *Tgfb1-3*, and *Foxp3* in *Dgka^-/-^z^f/f^-Foxp3^YFPCre/YFPCre^* and WT-*Foxp3^YFPCre/YFPCre^*Tregs. **J.** Overlaid histograms comparing expression of indicated molecules in *Dgka^-/-^z^f/f^-Foxp3^YFPCre/YFPCre^*and WT-*Foxp3^YFPCre/YFPCre^* mice. Data shown are representative of or pooled from at least three experiments. *, p < 0.05; **, p < 0.01; ***, p < 0.001; ****, p < 0.0001 determined by two-tail unpaired Student *t* test and pairwise Student *t* test for E.

αζDKO-Tregs displayed enhanced in vitro contact inhibition capability (Figure 4E), which is associated with their increased cell numbers after 72-hour incubation (Figure 4F). Moreover, αζDKO-Tregs proliferated more vigorously than WT-Tregs after in vitro TCR stimulation (Figure 4G). Thus, αζDKO-Tregs showed strong contact inhibition at least partially as a result of enhanced expansion. At present, it is unclear if individual αζDKO Treg exhibits stronger inhibition activity than a WT Treg.

αζDKO-Tregs manifested abnormal expression of many Treg-associated genes (Figure 4H, 4I). *Ctla4*, *Pdcd1* (encoding PD-1), *Tigit*, *Ezh2*, *Havcr2* (encoding Tim3), *Icos*, *Gzmb*, and *Klrg1* mRNA levels were increased, while *Foxp3*, *Il2ra*, *Il2rb*, *Il7ra*, *Entpd1* (encoding CD39), and *Nrp1* mRNA levels were decreased. At the protein level, CTLA-4, CD73, TIGIT, CD69, ICOS, GITR, and PD-1 were increased, and KLRG1 was not changed, but CD25 was decreased in either αζDKO cTregs, eTregs, or both (Figure 4J). Increased CTLA-4, TIGIT, GzmB, ICOS, and CD73 as well as *Tgfb3* and *Il10* suggested that certain aspects of Treg function might be enhanced while decreased C*d39*, *Il2ra*, *Il2rb*, *Il7r*, and *Ezh2* might negatively affect Treg function, homeostasis, and/or stability (40, 41). The decreased *Foxp3* mRNA, although only 10%, and *Nrp1* mRNA levels might also contribute to or reflect reduced stability. Increased CD69 expression also supported enhanced DAG signaling. The drastically upregulated PD-1 in both cTregs and eTregs may have inhibited Treg function and raised the possibility of exhaustion. Some of these changes were also observed in *Dgka^-/-^z^f/f^-Fopx3^YFPCre/+^* female mice (Supplemental Figure S6E), suggesting that DGKαζ double deficiency intrinsically influenced Treg properties.

### Altered Treg effector lineages and proinflammatory functions of **αζ**DKO Tregs

Tregs differentiate to multiple effector lineages to fulfil specific suppressive functions in different settings. T-bet^+^ Treg1, Gata3^+^, IRF4^+^, and Batf^+^ Treg2, RORγt^+^ and Stat3^+^ Treg17, and Bcl6^+^ Tfr-cells selectively suppress Th1, Th2, Th17, and Tfh cell/GC responses (19, 42–48). As shown in Figure 5A, αζDKO-Tregs upregulated TFs associated with Th2 (*Maf* and *Batf*), Th17 (*Rora*, *Rorc*, *Maf*, *JunD*, *Pou2af1* (encoding Bob1/BOF1)), and Tfh (*Bcl6*, *Ascl2*, *Batf*, *Pou2af1*, and *Maf*) but not Th1 lineages (*Tbx21*). Increased RORγt and Bcl6 were further confirmed at the protein level by intracellular staining (Figure 5B, 5C, Supplemental Figure S7A). Although *Gata3* mRNA was not increased, its protein was upregulated in αζDKO-Tregs, suggesting posttranscriptional regulation. Moreover, αζDKO-Tregs were enriched for Th2-, Th17-, and Tfh-like signatures (Supplemental Figure S7B, S7C). Thus, differentiation of multiple Treg effector lineages, except Th1-like Tregs, were enhanced in Treg-αζDKO mice.

**Figure 5.**
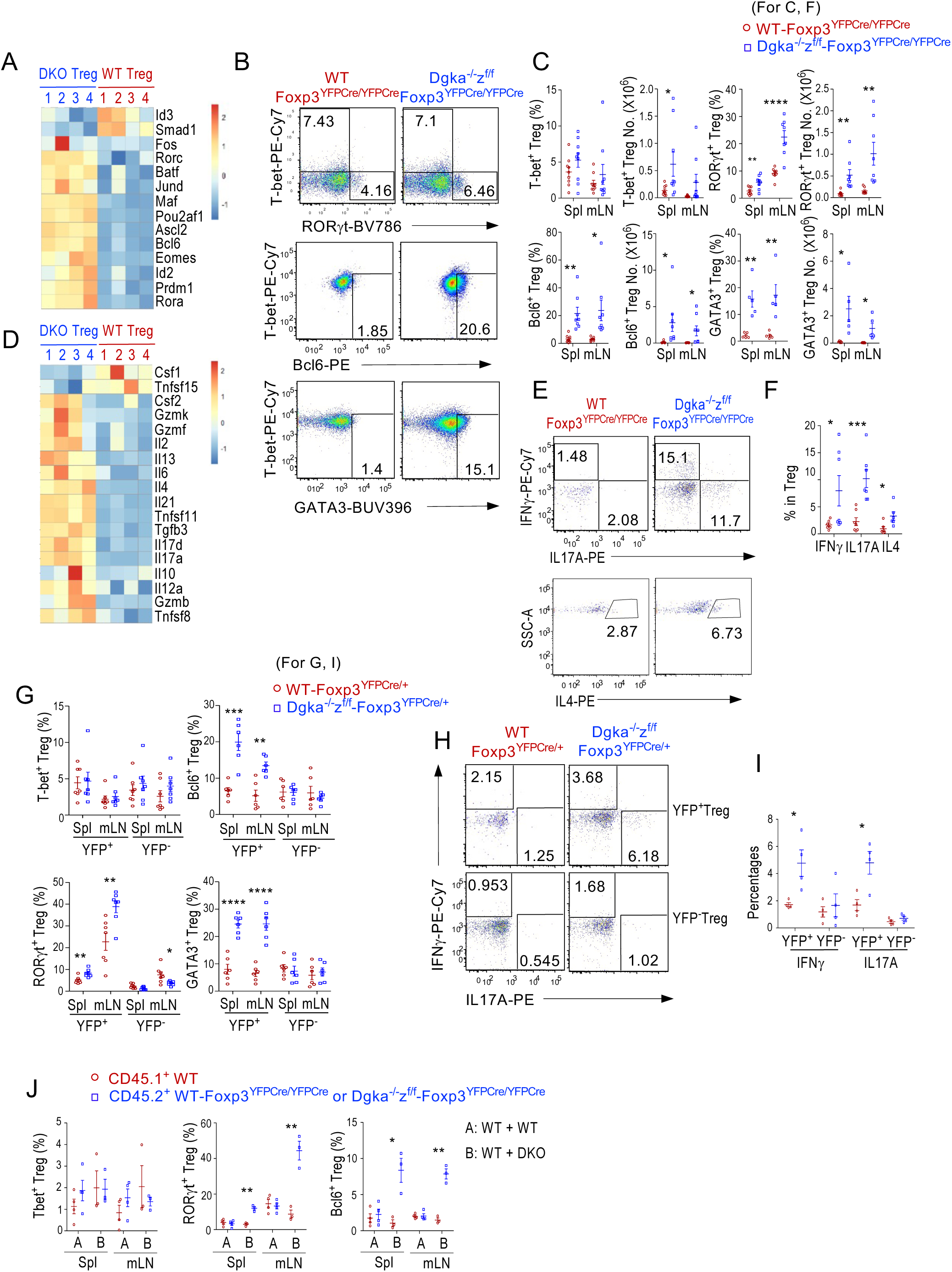
Increased effector lineages and gain-of-proinflammatory properties of αζDKO Tregs. **A–F.** Analyses of WT-*Foxp3^YFPCre/YFPCre^* and *Dgka^-/-^z^f/f^-Foxp3^YFPCre/YFPCre^*Tregs. **A.** Heatmap showing DE of TFs in Tregs by RNA sequencing. **B.** Intracellular staining of RORγt, T-bet, Bcl6, and GATA3 in splenic Tregs. **C.** Percentages and numbers (mean ± SEM) of Treg subsets. **D.** Heatmap showing DE of cytokines in Tregs by RNA sequencing. **E.** Intracellular staining of cytokines in Tregs after ex vivo PMA plus ionomycin stimulation in the present GolgiPlug for 5 hours. **F.** Percentages (mean ± SEM) of IFNγ^+^, IL17A^+^, and IL4^+^ Tregs. **G–I.** Analyses of female WT*-Foxp3^YFPCre/+^* and *Dgka^-/-^z^f/f^ -Foxp3^YFPCre/+^* Tregs. **G.** Percentages (mean ± SEM) of Treg effector sublineages in YFP^+^ and YFP^-^ Tregs. **H–I.** IL17A and IFNγ expression in Tregs after ex vivo PMA plus ionomycin stimulation in the present GolgiPlug for 5 hours. **H.** Representative FACS plots of Tregs. **I.** Scatter plots show mean ± SEM of IFNγ^+^ and IL17A^+^ in YFP^+^ and YFP^-^ Tregs. **J.** Analyses of mixed BM chimeric mice described in Figure 2N. Scatter plots show mean ± SEM of Treg sublineage percentages in CD45.1^+^ WT (red circle) and CD45.2^+^ WT or αζDKO Tregs (blue square). Data shown in B–J are representative of or pooled from at least three experiments. *, p < 0.05; **, p < 0.01; ***, p < 0.001; ****, p < 0.0001 determined by two-tail unpaired Student *t* test.

Because Tregs express TCRs with relatively high affinity to self-antigens, suppression of proinflammatory cytokine expression in Tregs is important for preventing Tregs from potential self-damage (1). Strikingly, αζDKO-Tregs expressed increased levels of *Il2*, Th2 cytokines *Il4* and *Il13*, Th17 cytokines *Il17a* and *Il17d*, Tfh cytokine *Il21*, proinflammatory cytokines *Il6* and *Csf2* (GM-CSF), and cytotoxicity-associated effectors *Gzmb*, *Gzmf*, and *Gzmk* (Figure 5D). Increased expression of IL17A and IL4 was further confirmed by intracellular staining (Figure 5E, 5F). Although T-bet and *Ifng* mRNA were not increased, IFNγ protein was increased, suggesting posttranscriptional regulation of IFNγ by DGKαζ.

The increases of GATA3^+^, RORγt^+^, and Bcl6^+^ sublineages and elevated IL17A and IFNγ production in αζDKO-Tregs were also observed in the YFP^+^ Tregs in female *Dgka^-/-^z^f/f^-Foxp3^YFPCre/+^* mice (Figure 5G–5I) as well as in the CD45.2^+^ αζDKO-Treg in mixed BM chimeric mice (Figure 5J).

Together, these data revealed that DGKα and ζ intrinsically suppress Treg effector lineage differentiation and also inhibit expression of multiple cytokines including Th effector cytokines IL-4, IL-17, and IFNγ in Tregs at transcriptional and posttranscriptional levels.

### Deregulated effector functions of CD4^+^Foxp3^-^ and CD8^+^ T cells in Treg-**αζ**DKO mice

Abnormal Treg functions could lead to deregulation of CD4^+^Foxp3^-^ conventional T cells (Tcons) and CD8^+^ T-cells. In Treg-αζDKO mice, CD4^+^Foxp3^-^ Tcons and CD8^+^ T-cells were increased in number in spleen and LNs (Figure 6A) at least partially as a result of enhanced proliferation (Figure 6B, 6C), although their percentages were decreased owing to disproportional expansion of B-cells, as detailed in Figure 7F. They contained decreased naïve but increased CD44^+^CD62L^+^ central memory (CM) and/or CD44^+^CD62L^-^ effector memory (EM) cells (Figure 6D, 6E), accompanying increased T-bet^+^ and IFNγ-producing CD8^+^ T-cells (Figure 6F–6H), suggesting impaired Treg suppression on CD8^+^ T-cells in these mice.

**Figure 6.**
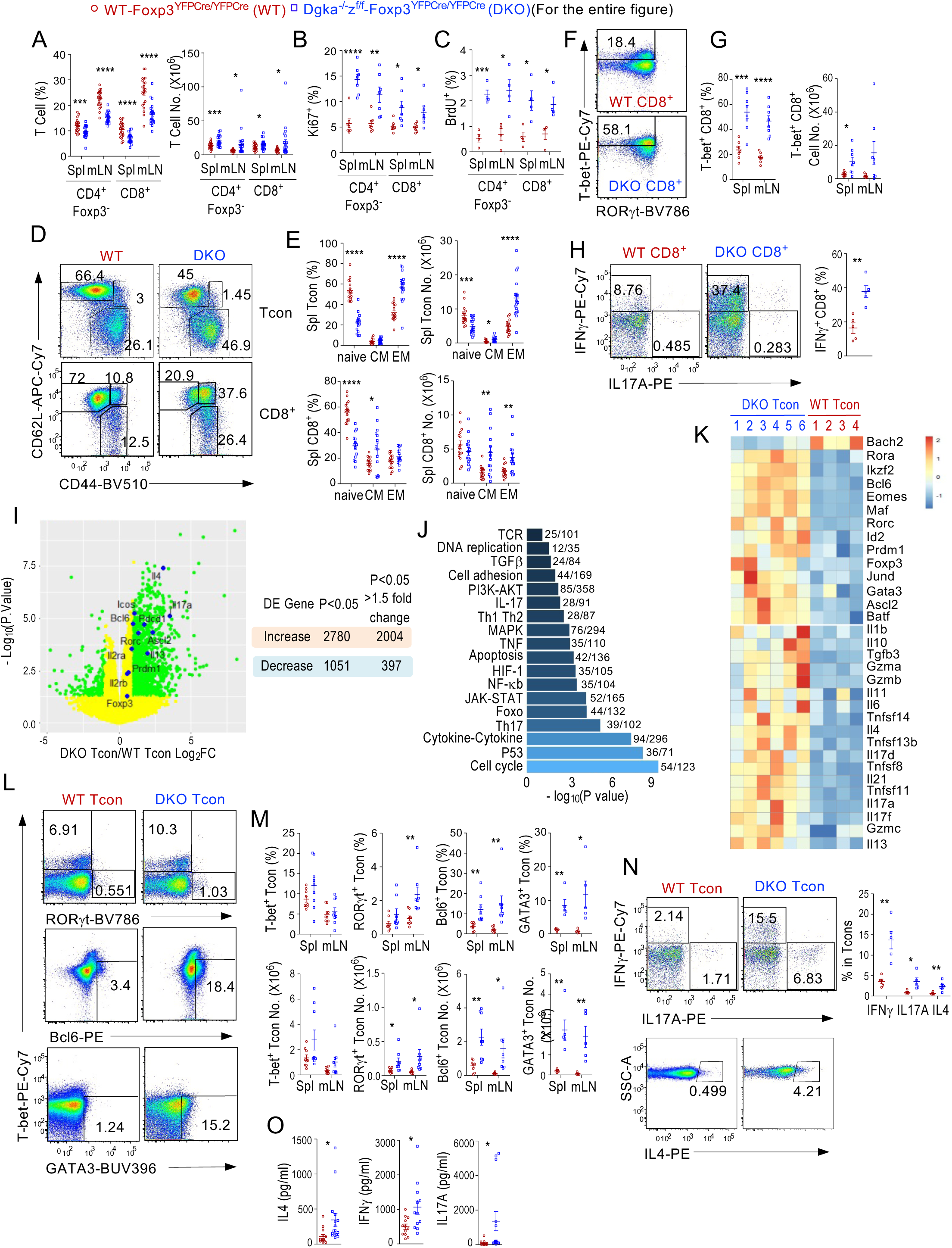
Enhanced effector function of CD4^+^Foxp3^-^ Tcon and CD8 T cells in Treg-αζDKO mice. *Dgka^-/-^z^f/f^-Foxp3^YFPCre/YFPCre^* and WT*-Foxp3^YFPCre/YFPCre^* control mice were analyzed. **A.** CD4^+^Foxp3^-^ Tcon and CD8^+^ T cell percentages and numbers in the spleen and mLNs. **B.** Ki67^+^ cells in Tcon and CD8 T cells. **C.** BrdU^+^ cells in Tcon and CD8 T cells. **D.** Representative FACS plots showing CD44 and CD62L expression in splenic CD4^+^Foxp3^-^ Tcon and CD8^+^ T cells. **E.** Naïve, CM, and EM percentages and numbers of splenic CD4^+^Foxp3^-^ Tcon and CD8 T cells. **F.** Representative FACS plots showing intracellular T-bet and RORγt staining in LN CD8 T cells. **G.** Percentages and numbers of T-bet^+^ CD8 T cells. **H.** Intracellular IFNγ and IL17A staining in CD8 T cells after PMA and ionomycin stimulation. Scatter plot represents mean ± SEM of IFNγ^+^ CD8 T cells from 6–12-month-old mice. **I.** Volcano plot comparing mRNA expression in WT and Treg-αζDKO CD4^+^Foxp3^-^ Tcons after RNA-seq analysis. Genes colored in green are differentially expressed with greater than 1.5-fold differences between WT and αζDKO Tregs (p < 0.05). Table shows numbers of DE genes (p < 0.05). **J.** Top enriched KEGG pathways between WT and Treg-αζDKO CD4^+^Foxp3^-^ Tcons. **K.** Heatmap showing mRNA levels of TFs and cytokines that were differentially expressed between WT and Treg-αζDKO CD4^+^Foxp3^-^ Tcons (p < 0.05). **L.** T-bet, RORγt, Bcl6, and GATA3 proteins in splenic Tcons detected by intracellular staining. **M.** Percentages and numbers of T-bet^+^, RORγt^+^, Bcl6^+^, and GATA3^+^ cells in Tcons. **N.** IFNγ, IL17A, and IL4 protein levels in WT and Treg-αζDKO CD4^+^Foxp3^-^ Tcons detected by intracellular staining after PMA and ionomycin stimulation for 5 hours. **O.** Serum cytokine levels in 5–12-month-old WT and Treg-αζDKO mice. Data shown are representative of or pooled from 4–22 experiments. *, p < 0.05; **, p < 0.01; ***, p < 0.001; ****, p < 0.0001 determined by two-tail unpaired Student *t* test.

**Figure 7.**
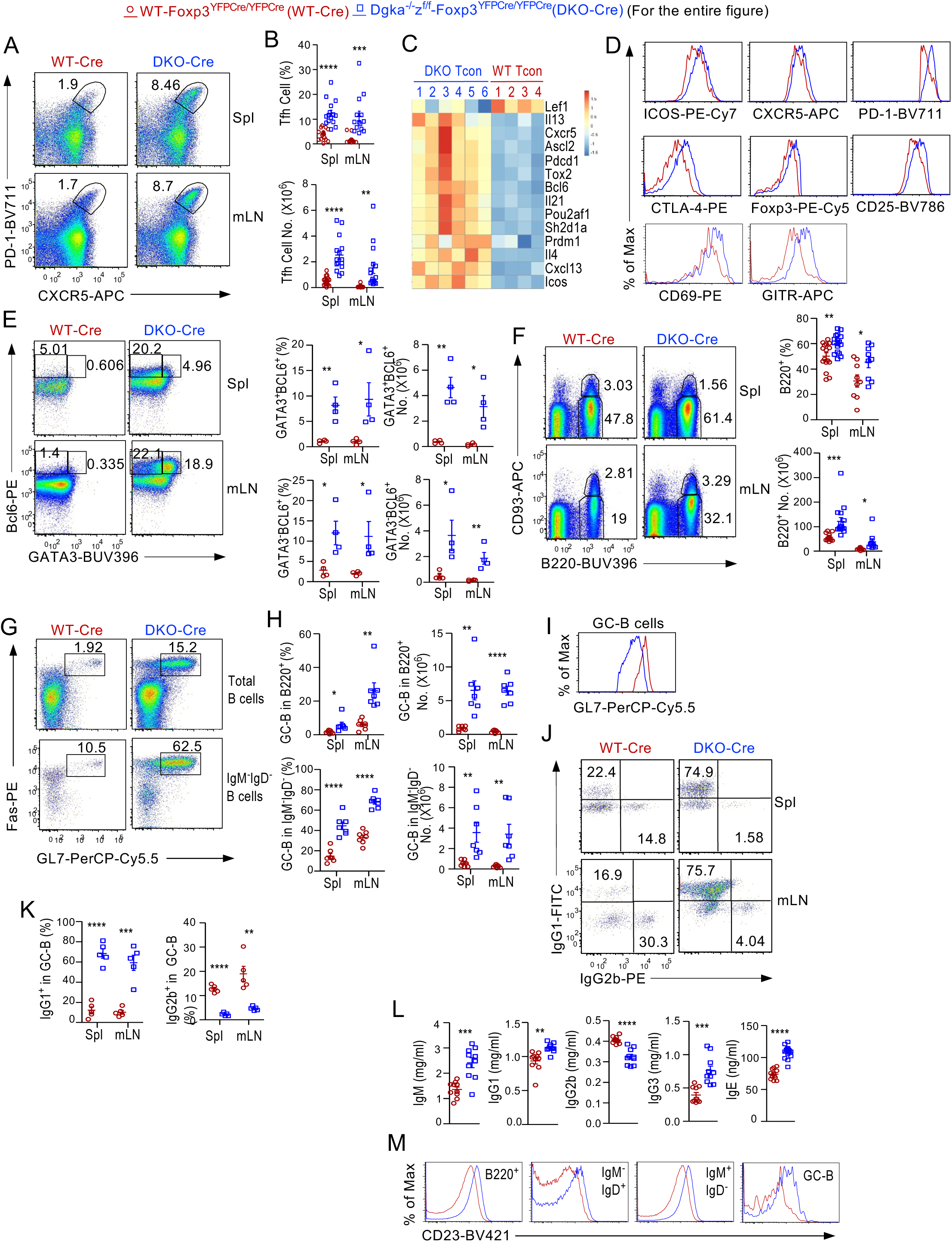
Enhanced Tfh/GC-B cell responses in *Dgka^-/-^z^f/f^-Foxp3^YFPCre/YFPCre^*mice. Splenocytes and LN cells from *Dgka^-/-^z^f/f^-Foxp3^YFPCre/YFPCre^*and WT-*Foxp3^YFPCre/YFPCre^* mice were analyzed. **A.** CXCR5 and PD-1 expression in CD4^+^Foxp3^-^ Tcons. **B.** Percentages and numbers of CXCR5^+^PD-1^+^ Tfh-cells in 2–14-month-old mice. **C.** Heatmap shows DE of key Tfh/Tfr genes from transcriptomic analyses of Tcons described in Figure 6I. **D.** Overlaid histograms show expression-indicated molecules in Tfh cells. **E.** Representative FACS plots show GATA3 and Bcl6 expression in Tcons from a pair of 7-month-old mice. Scatter plots show mean ± SEM of GATA3^+^Bcl6^+^ and GATA3^-^Bcl6^+^ Tfh cells. **F.** B220 and CD93 staining of splenocytes and LN cells. Scatter plots show mean ± SEM of B220^+^CD93^-^ mature B-cell percentages and numbers. **G.** GL7 and Fas expression in total B220^+^ and in IgM^-^IgD^-^B220^+^ cells. **H.** Scatter plots show mean ± SEM of GC-B cell percentages and numbers in 5–14-month-old mice. **I.** Overlaid histogram shows GL7 expression in GC-B cells. **J.** Intracellular IgG1 and IgG2b staining in GC-B cells. **K.** IgG1^+^ and IgG2b^+^ percentages in GC-B cells. **L.** Serum IgM, IgG1, IgG2b, IgG3, and IgE concentrations. **M.** CD23 expression in B cell populations. Data shown are representative of or pooled from at least four experiments. *, p < 0.05; **, p < 0.01; ***, p < 0.001; ****, p < 0.0001 determined by two-tail unpaired Student *t* test.

To facilitate the understanding of abnormalities in CD4^+^ Tcons, we performed transcriptomic analyses of CD4^+^Foxp3YFP^-^ Tcons from WT*-Foxp3^YFPCre/YFPCre^*and *Dgka^-/-^z^f/f^-Foxp3^YFPCre/YFPCre^* mice. Because CD4^+^Foxp3YFP^-^ Tcons from *Dgka^-/-^z^f/f^-Foxp3^YFPCre/YFPCre^* mice should contain WT (more accurately, DGKα-deficient) Tcons and DGKαζ double-deficient ex-Foxp3 cells (exTregs), we reasoned that such analyses would be informative for revealing abnormalities of Tcons resulting from impairment of certain aspects of Treg functions and potential gained properties of exTregs arising from absence of both DGKα and ζ.

Treg-αζDKO CD4^+^Foxp3^-^ Tcon transcriptome displayed obvious differences from control Tcon’s. They had 2,780 upregulated and 1,051 downregulated genes (p < 0.05, Supplemental Table 3) including 2,004 and 397 of them respectively with differences greater than 1.5 folds (Figure 6I). One hundred and thirty-four KEGG pathways, with striking similarities to those of Tregs, were enriched in αζDKO Tcons (Figure 6J, Supplemental Table 4). Cell cycle and DNA replication were also among the top enriched pathways with many cell cycle-promoting molecules upregulated (Supplemental Figure S8A), further supporting dysregulated Tcon expansion.

Treg-αζDKO CD4^+^Foxp3^-^ Tcons were also enriched with Th1, Th2, Th17, and Tfh signatures (Figure 6J, 6K, Supplemental Figure S8B–S8D). mRNA levels of multiple TFs associated with effector differentiation (*Eomes*, *Prdm1* [encoding Blimp1], and *Klrg1*), Th2 differentiation (*Gata3*, *cMaf, Batf*), Th17 differentiation (*Rora*, *Rorc*, *cMaf*, and *Jund*), and Tfh differentiation (*Bcl6*, *Batf*, *Ascl2*, and *Pou2af1*) were increased (Figure 6K). Expression of *Bach2*, which inhibits Tfh and Th17 differentiation to promote Treg stability (49), was decreased. Consistently, RORγt^+^, Bcl6^+^, GATA3^+^, and, to a lesser extent, T-bet^+^ cells within Treg-αζDKO CD4^+^Foxp3^-^ Tcons were increased (Figure 6L, 6M), accompanying increased mRNA of many Th2 (*Il4* and *Il13*), Th17 (*Il17a/d/f*), and Tfh (*Il21*) cytokines (Figure 6K) and IL4, IL17A, and IFNγ proteins (Figure 6N). In some old Treg-DKO mice, serum IL4, IFNγ, and IL17A levels were elevated (Figure 6O). CD4^+^Foxp3^-^ T cells from *Dgka^-/-^z^f/f^-Foxp3^YFPCre/YFPCre^*expressed altered cell surface markers. They had upregulated CTLA-4, ICOS, PD-1, CD73, TIGIT, and several other molecules (Supplemental Figure S8E). Thus, Treg-αζDKO CD4 Tcons, remarkably similar to Tregs, displayed enhanced Th2/17 and Tfh differentiation.

Together, these data indicated that deficiency of both DGKα and ζ in Tregs led to enhanced proliferation and effector functions of both CD4^+^Foxp3^-^ Tcons and CD8^+^ T-cells. Such abnormalities might play a role in the overall inflammatory and autoimmune status in Treg-αζDKO mice and contribute to the phenotypes we observed in these mice.

### Enhanced Tfh- and Tfh2/13-skewed GC responses in Treg-**αζ**DKO mice

Tfh-cells are crucial for GC responses and humoral immunity. In *Dgka^-/-^z^f/f^-Foxp3^YFPCre/YFPCre^* mice, Foxp3^-^CD4^+^CXCR5^+^PD-1^+^ Tfh-cell percentages and numbers were drastically increased (Figure 7A, 7B), which was consistent with the enrichment of the Tfh pathway and increased expression of Tfh TFs *Bcl6*, *Ascl2*, and *Pou2af1* (50–52) and effector molecules *Il21*, *Il4*, and *Il13* in CD4^+^Foxp3^-^ Tcons (Figure 7C, 6K–6M, Supplemental Figure S8D). Treg-αζDKO Tfh cells upregulated Tfh-promoting molecule ICOS and CXCR5; T-cell activation markers CD69 and PD-1; and Treg-associated molecules CTLA-4, CD25, GITR, and Foxp3 (Figure 7D, Supplemental Figure S9A), with both GATA3^+^Bcl6^+^ IL4/IL13-expressing Tfh2/13-cells (53) and GATA3^-^Bcl6^+^ Tfh-cells increasing (Figure 7E) and elevated GATA3 levels in Tfh cells (Supplemental Figure S9B).

Consistent with increased Tfh-cells, *Dgka^-/-^z^f/f^-Foxp3^YFPCre/YFPCre^*mice had increased B220^+^ B-cells, mostly as a result of increases of IgM^-^IgD^+^ and IgM^-^IgD^-^ cells (Figure 7F, Supplemental Figure S9C) and increased GL7^+^Fas^+^ GC-B-cells and GL7^-^Fas^+^ activating B-cells within total B220^+^ or IgM^-^IgD^-^B220^+^ cells (Figure 7G, 7H, Supplemental Figure S9D). GC-B cells from Treg-αζDKO mice showed decreased GL7 levels, likely reflecting overactivation (Figure 7I), enhanced proliferation but similar survival (Supplemental Figure S9E, S9F), and markedly increased IgG1^+^ but reduced IgG2b^+^ ratios (Figure 7J, 7K). Additionally, IgM^-^IgD^-^ FAS^+^GL-7^-^ and IgM^-^IgD^-^FAS^-^GL-7^-^ non-GC B cell numbers were increased and both populations contained increased IgG1 but decreased IgG2b ratios (Supplemental Figure S9G), Consistently, CD4^+^Foxp3^-^ CXCR5^-^PD-1^+^ T peripheral helper (Tph) cells, which promote extrafollicular B cell antibody responses and contribute to autoimmunity (54–57), were increased in both percentages and numbers (Supplemental Figure S9H), associated with increased proliferation but not survival (Supplemental Figure S9I) and upregulated ICOS levels (Supplemental Figure S9J). Treg-αζDKO mice contained increased serum IgM, IgG1, IgE, and IgG3 levels but decreased IgG2b levels (Figure 7L), accompanying obviously increased CD23 (the low affinity receptor for IgE) in various B-cells (Figure 7M), which is consistent with the ability of serum IgE to upregulate CD23 (58).

Together, DGKαζ deficiency in Tregs led to enhanced Tfh and GC-B cell as well as non-GC B cell responses with prominent polarization to the Tfh2/13 lineage and IgG1/IgE responses, which provided mechanistic explanation of development of autoantibodies and lupus-like diseases and the IgG1-predominant autoimmunity in Treg-αζDKO mice. The increased Tfh and GC-B cells could be caused by a potential impaired Treg/Tfr-cell mediated suppressive mechanism yet to be defined, by increased Treg/Tfr-cells to exTreg-Tfh cell conversion (as will be described later), by the inflammatory environment of dysregulated T helper cells and effector CD8 T cells, and/or by positive feedbacks or bystander activations between Tfh and GC B cells.

### Dysregulated Tfr-cells in Treg-**αζ**DKO mice

Tfr-cells suppress GC responses by inhibiting Ig class-switch and Tfh-cell function, and their differentiation is Bcl6-dependent (18, 19, 46). In *Dgka^-/-^z^f/f^-Foxp3^YFPCre/YFPCre^*mice, Foxp3^+^CD4^+^CXCR5^+^PD-1^+^ Tfr-cells were increased (Supplemental Figure S10A, S10B), at least as a result of enhanced proliferation but not survival (Supplemental Figure S10C, S10D). Within CXCR5^+^PD-1^+^ CD4^+^T cells, the Foxp3^+^ Tfr to Foxp3^-^ Tfh cell ratios were increased (Supplemental Figure S10E, S10F). αζDKO Tfr-cells upregulated CTLA-4, PD-1, CD69, and ICOS but downregulated CD25 and Nrp1 (Supplemental Figure S10G, S10H). Mature Tfr cells downregulate CD25 to avoid IL-2 signal mediated inhibition (59–61). Decreased CD25 in αζDKO Tregs and Tfr-cells could promote Tfr-cells differentiation, which might partially contribute to increased Tfr-cells. αζDKO Tfr-cells also upregulated Bcl6 (Supplemental Figure S10I), accompanying increased GATA3^+^Bcl6^+^ Tfr2/13-cells and GATA3^-^Bcl6^+^ Tfr-cells (Supplemental Figure S10I, S10J) and thus bias toward Tfr2/13-cells. Consistent with increased Tfr-cells, αζDKO Tregs were enriched in Tfr/Tfh-associated gene signatures (Supplemental Figure S10K), characterized by upregulation of positive regulators (*Bcl6*, *Cxcr5*, *Pdcd1*, *Ascl2*, *Batf*, *Icos*, *Pou2af1*, *Cxcr5*, and *Sh2d1a* [encoding the adaptor molecule SAP]) of Tfh-cell and/or Tfr-cell differentiation. Although it is unclear whether αζDKO Tfr-cells were impaired in suppressive function, they gained expression of Tfh-associated cytokine *Il21* as well as Th2-associated cytokines *Il4* and *Il13*, which might have contributed to the IgG1/IgE-predominant antibodies and autoimmunity in Treg-αζDKO mice. Of note, some Tfh cells upregulate Foxp3 in GCs (62). Of note, our data do not rule out that αζDKO Tfr-cells might contain some Tfh-derived Foxp3^+^ cells.

### Development of lupus-like diseases in female *Foxp3^YFPCre/+^* heterozygous *Dgka^-/-^z^f/f^-Foxp3^YFPCre/+^* mice

Female *Dgka^-/-^z^f/f^-Foxp3^YFPCre/+^* (DKO-Cre^het^) mice contained both *Dgka^-/-^* control (called WT for simplicity) and αζDKO-Tregs. Surprisingly, they also lost weight and manifested lymphoproliferative/autoimmune disorders, albert less severely than *Dgka^-/-^z^f/f^-Foxp3^YFPCre/YFPCre^*mice (Figure 8A–8F, Supplemental Figure S11A). They had increased B-cells (Figure 8G); GC B-cells within both total B220^+^ and B220^+^IgM^-^IgD^-^ DN B-cells (Figure 8H, 8I); serum IgM, IgG1, and IgE (Figure 8J); and IgG1^+^ in GC B-cells (Figure 8K) but decreased serum IgG2b and IgG3 and IgG2b^+^ cells. The increased CD23 in B-cells further supported elevated serum IgE (Supplemental Figure S11B). In DKO-Cre^het^ mice, CD4^+^Foxp3^-^ Tcons were decreased in percentages but not in numbers owing to disproportional increases of B-cells (Figure 8L). They contained increased effector but decreased naïve T-cells (Figure 8M) accompanying increased percentages of GATA3^+^, RORγt^+^, and Bcl6^+^ but not T-bet^+^ CD4^+^Foxp3^-^ Tcons (Figure 8N). They had increased Tfh-cells (Figure 8O) with elevated PD-1, ICOS, and CXCR5 (Supplemental Figure S11C). In DKO-Cre^het^ mice, YFP^+^ but not YFP^-^ Tfr-cells were increased (Figure 8P). The YFP^+^Tfr/Tfh cell ratios were increased but YFP^-^/Tfh cell ratios were decreased, although total Tfr/Tfh cell ratios were not obviously altered (Figure 8Q). Together, these data suggested that DGKα and ζ intrinsically inhibited Tfr-cell differentiation and that αζDKO Tregs/Tfr-cells and/or exTregs gained dominant functions that could overpower WT Treg/Tfr-cells to cause autoimmune diseases.

**Figure 8.**
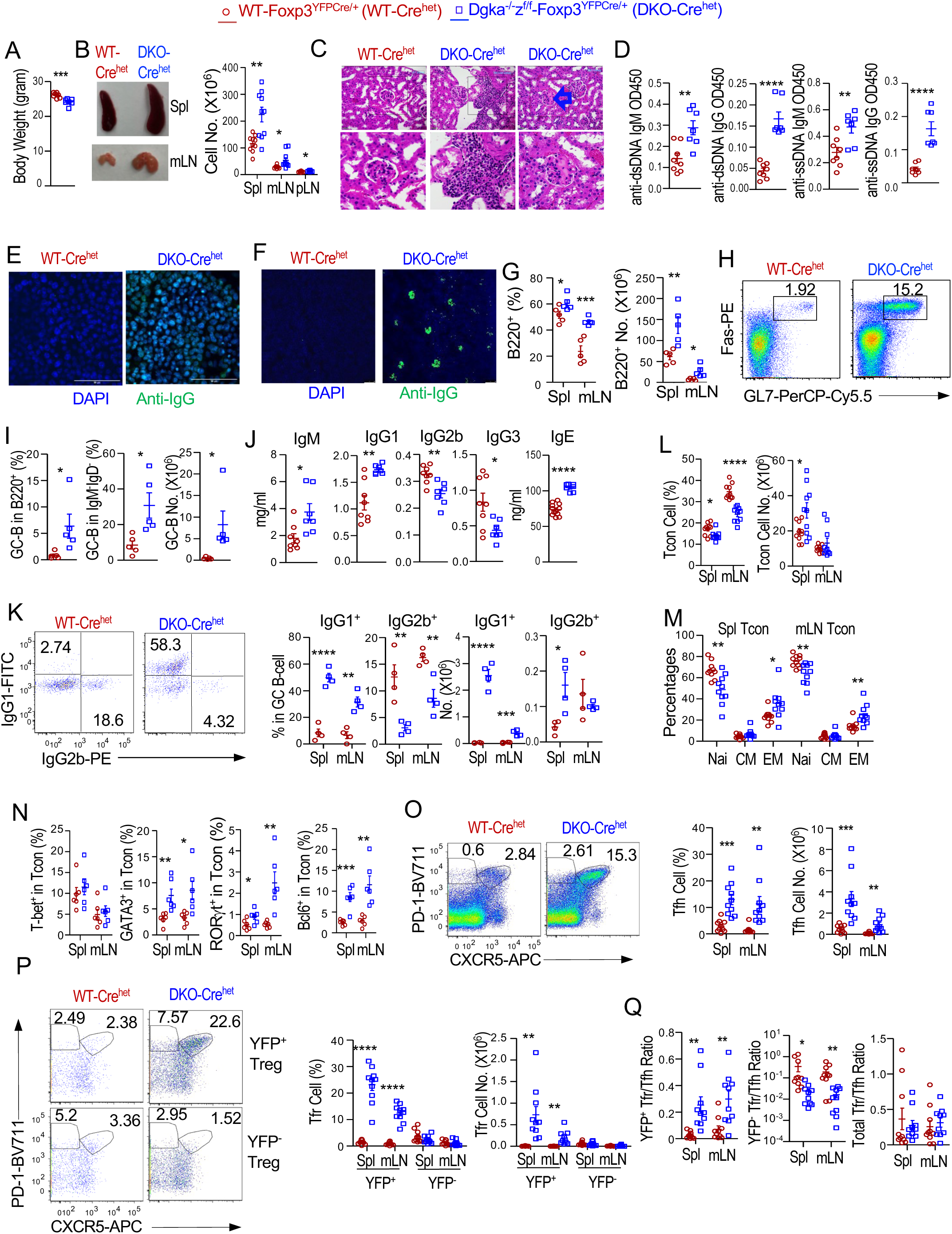
Development of autoimmune diseases and deregulated Tfh/Tfr cell and GC-responses in female *Dgka^-/-^z^f/f^-Foxp3^YFPCre/+^* mice. Three–nine-month-old female *Dgka^-/-^z^f/f^-Foxp3^YFPCre/+^* (DKO-Cre^het^) and WT*-Foxp3^YFPCre/+^*(WT-Cre^het^) mice were analyzed. **A.** Body weights. **B.** Representative picture of spleen and mLNs, total cell numbers in the indicated organs. **C.** H&E staining of kidney thin sections. Bottom low shows higher magnification. **D.** Seral anti-dsDNA and ssDNA autoantibodies. **E.** Antinuclear antibodies. **F.** IgG deposition in the kidney. **G.** Total B cell percentages and numbers. **H.** Fas and GL7 staining in splenic B220^+^ cells. **I.** GC-B cell percentages in splenic B220^+^ and B220^+^IgM^-^IgD^-^ (DN) B cells and GC-B cell numbers in B220^+^ B cells. **J.** Seral Ig levels. **K.** IgG1^+^ and IgG2b^+^ cells in GC-B cells. **L.** CD4^+^Foxp3^-^ Tcon percentages and numbers. **M.** Naïve and effector cell percentages in Tcons. **N.** Percentages of T-bet^+^, GATA3^+^, RORγt^+^, and Bcl6^+^ cells in Tcons. **O.** Assessment of Tfh cells. Representative FACS plots show gating of CXCR5^+^PD-1^+^ Tfh and CXCR5^-^PD-1^+^ Tph cells in splenic Tcons. Scatter plots show Tfh percentages and numbers. **P.** Assessment of Tfr cells. Representative FACS plots show gating of CXCR5^+^PD-1^+^ Tfr cells in splenic YFP^+^ and YFP^-^ Foxp3^+^ Tregs. Scatter plots show Tfr percentages and numbers. **Q.** Scatter plots show Tfr/Tfh cell ratios. Data shown are representative of or pooled from 5–10 experiments except four experiments for Figure 8K. *, p < 0.05; **, p < 0.01; ***, p < 0.001; ****, p < 0.0001 determined by two-tail pair-wise (Figure 8A, data with lines connecting WT-Cre^het^ and αζDKO-Cre^het^ mice) or unpaired Student *t* test.

### DGK**αζ** deficiency conferred CD28-independent Treg, Tfr-cell, and exTreg-Tfh-cell development and homeostasis and dysregulated GC responses and autoimmune diseases

CD28 costimulatory signal promotes Ras-Erk1/2 and PI3K/mTOR signaling and is critical for Treg and Tfr development and homeostasis and for Tfh differentiation to promote GC responses and humoral immunity (63–65). Enhanced mTORC1/2 and Erk1/2 activation in αζDKO-Tregs prompted us to examine whether Treg-αζDKO relieved their dependence on CD28. *Foxp3^YFPCre/YFPCre^-CD28^-/-^* (28KO-Cre) mice had very few Tregs; *Dgka^-/-^z^f/f^-Foxp3^YFPCre/YFPCre^-CD28^-/-^* (TKO-Cre) Tregs had increased slightly in the thymus and obviously in the periphery (Figure 9A), accompanying increased e/cTreg ratios (Figure 9B).

**Figure 9.**
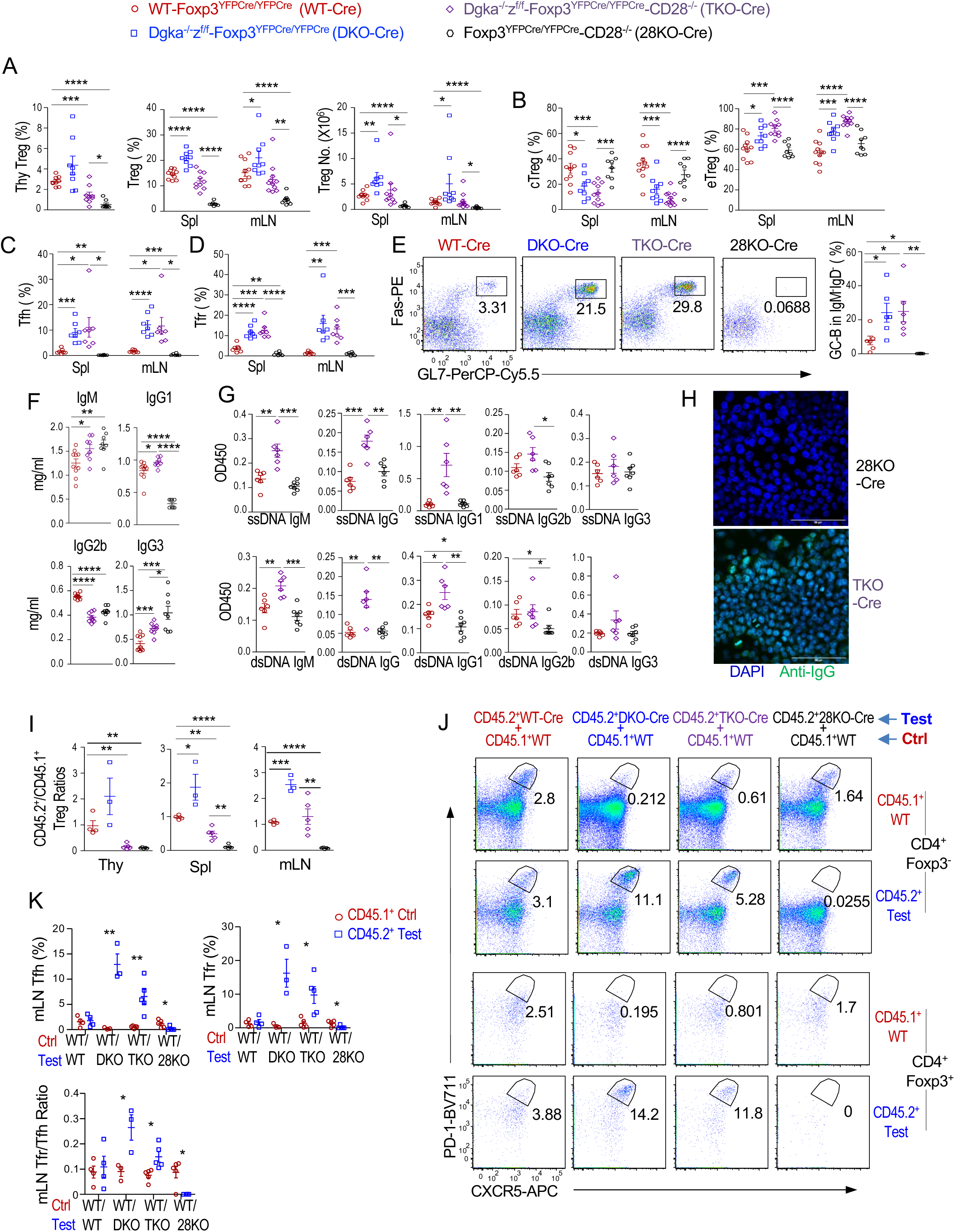
Treg-specific DGKαζ deficiency conferred CD28-independent Treg development/homeostasis and GC responses and accelerated Treg-to-exTreg/exTreg-Tfh conversion. **A–H.** Analyses of WT*-Foxp3^YFPCre/YFPCre^*, *Dgka^-/-^z^f/f^-Foxp3^YFPCre/YFPCre^* (DKO), *Dgka^-/-^z^f/f^-Foxp3^YFPCre/YFPCre^-CD28^-/-^* (TKO), and *Foxp3^YFPCre/YFPCre^-CD28^-/-^* (CD28KO). **A.** Treg percentages and numbers (mean ± SEM) in the thymus, spleen, and mLNs. **B.** Mean ± SEM of cTreg and eTreg percentages. **C.** Tfh-cell percentages. **D.** Tfr-cell percentages. **E.** Representative FACS plots showing Fas and GL7 staining in live gated splenic B220^+^IgM^-^IgD^-^ (DN) B cells. Scatter plots show mean ± SEM of GC-B cell percentages. **F.** Seral antibody levels. **G.** Seral anti-ssDNA and dsDNA autoantibody levels. **H.** Seral antinuclear antibodies in *CD28^-/-^* and TKO mice. **I–K.** Analyses of mixed BM chimeric mice reconstituted with a mixture of BM cells of CD45.1^+^ WT BM cells with either CD45.2^+^ WT*-Foxp3^YFPCre/YFPCre^*, *Dgka^-/-^z^f/f^-Foxp3^YFPCre/YFPCre^*(DKO), *Dgka^-/-^z^f/f^-Foxp3^YFPCre/YFPCre^-CD28^-/-^*(TKO), or *Foxp3^YFPCre/YFPCre^-CD28^-/-^* (28KO) BM cells. **I.** Ratios of CD45.2^+^ test Treg percentages in CD4^+^ T cells/CD45.1^+^ WT Treg percentage in CD4^+^ T cells in individual chimeric mice. **J.** Representative FACS plots showing PD-1 and CXCR5 staining in live gated mLN CD45.1^+^ control and CD45.2^+^ test CD4^+^Foxp3^-^ Tcons and CD4^+^Foxp3^+^ Tregs. **K.** Percentages of Tfh and Tfr cells as well as Tfr/Tfh ratios of CD45.1^+^ and CD45.2^+^ origins in individual chimeric mice. Data shown are representative of or pooled from 6–11 experiments for A–H and 3–5 experiments for I–K. *, p < 0.05; **, p < 0.01; ***, p < 0.001; ****, p < 0.0001 by two-tailed unpaired Student t-test.

CD28KO-Cre mice were deficient in Tfh/Tfr-cells (Figure 9C, 9D) and GC B-cells (Figure 9E). They contained elevated serum IgM and IgG3 but reduced IgG1 and IgG2b (Figure 9F). However, most of these phenotypes, with the exception of IgG2b, were reversed in TKO-Cre mice. TKO-Cre mice even had more Tfh, Tfr, and GC B-cells as well as increased Tfr/Tfh cell ratios than WT mice (Figure 9C – 9E, Supplemental Figure S12A). Similar to αζDKO-Cre mice, TKO-Cre mice also developed IgG1-dominant autoantibodies and multiorgan autoimmune diseases (Figure 9G, 9H, Supplemental Figure S12B). Thus, Treg-αζDKO not only completely or partially reversed defects in Tregs, Tfr, and GC B-cells and humoral immunity caused by CD28 deficiency but also triggered IgG1-predominant autoimmunity independent of CD28.

In mixed BM chimeric mice reconstituted with a mixture of BM cells of CD45.1^+^ WT BM cells with either CD45.2^+^ WT-Cre, αζDKO-Cre, TKO-Cre, or 28KO-Cre BM cells, TKO-Cre Tregs were also increased, especially in the spleen and LNs (Figure 9I). Thus, DGKαζ activities intrinsically enforce CD28 dependence for Treg development/homeostasis and cTreg-to-eTreg differentiation.

Highly interestingly, although virtually no Tfr and Tfh-cells developed from 28KO-Cre BM cells, TKO-Cre-derived CD4^+^Foxp3^+^ Tregs and CD4^+^YFP^-^Foxp3^-^ Tcons contained increased Tfr and Tfh-cells even compared with WT-Cre controls in the mixed BM chimeric mice (Figure 9J – 9K). The Tfr/Tfh ratios were also increased within TKO derived CD4^+^ T cells (Figure 9K). Because the canonical TKO-Cre BM cell-derived CD4^+^Foxp3YFP^-^ Tcons were CD28 deficient but expressed WT DGKζ, and should be defective in Tfh-/Tfr-cell differentiation, these TKO-derived Tfh-cells should have converted from Tregs/Tfr-cells and deficient of DGKζ as well as DGKα, and αζDKO in Tregs should have conferred CD28-independent Tfr/Tfh differentiation. However, although it is unlikely, these data do not rule out that DGKα deficiency in Tcons due to germline deficiency in the context of DGKαζ double deficiency in Tregs could play a role in promoting Tfh cell differentiation.

### Accelerated conversion of **αζ**DKO Treg to exTregs and exTreg-Tfh cells and pathogenicity of **αζ**DKO Tregs/exTregs in IgG1-predominant GC B-cell responses and autoimmunity

ExTregs can be pathogenic owing to their expression of TCRs with relatively high affinities to self-antigens. If increased conversion of Tregs to exTregs occurred in *Dgka^-/-^z^f/f^-Foxp3^YFPCre/YFPCre^* mice, it should blend the CD4^+^Foxp3YFP^-^ T-cell population with exTregs that might express residue Treg signature genes. Among the 3,831 and 4,563 DEGs in *Dgka^-/-^z^f/f^-Foxp3^YFPCre/YFPCre^* CD4^+^YFP^-^ Tcons and CD4^+^YFP^+^ Tregs compared with their WT controls, 2,054 of them were differentially expressed in both Tregs and Tcons (Figure 10A, Supplemental Table 5). Within these 2,054 genes, 1,483 and 497 genes were concordantly increased or decreased in both Treg-αζDKO Tregs and Tcons, respectively (Figure 10B). Eleven genes with unclear functions in Tregs were increased in αζDKO Tregs but decreased in αζDKO Tcons. Sixty-three genes, including several Treg signature genes *Il2ra*, *Il2rb*, and, particularly, *Foxp3* were discordantly increased in Treg-αζDKO Tcons but decreased in Treg-αζDKO Tregs (Figure 10C). These data suggested that some Treg-αζDKO Tcons contained residual Treg-associated genes and were likely blended with exTregs. Consistent with this notion, we detected Cre-mediated recombination of *Dgkz^f/f^* alleles in sorted CD4^+^Foxp3YFP^-^Tcons from *Dgka^-/-^z^f/f^-Foxp3^YFPCre/YFPCre^* mice and female *Dgka^-/-^z^f/f^-Foxp3^YFPCre/+^*mice but not from *Dgka^+/+^z^f/f^-Foxp3^YFPCre/YFPCre^* mice (Supplemental Figure S13A), suggesting increased Treg-to-exTreg conversion in Treg-specific DGKα and ζ double but not DGKζ single deficient mice.

**Figure 10.**
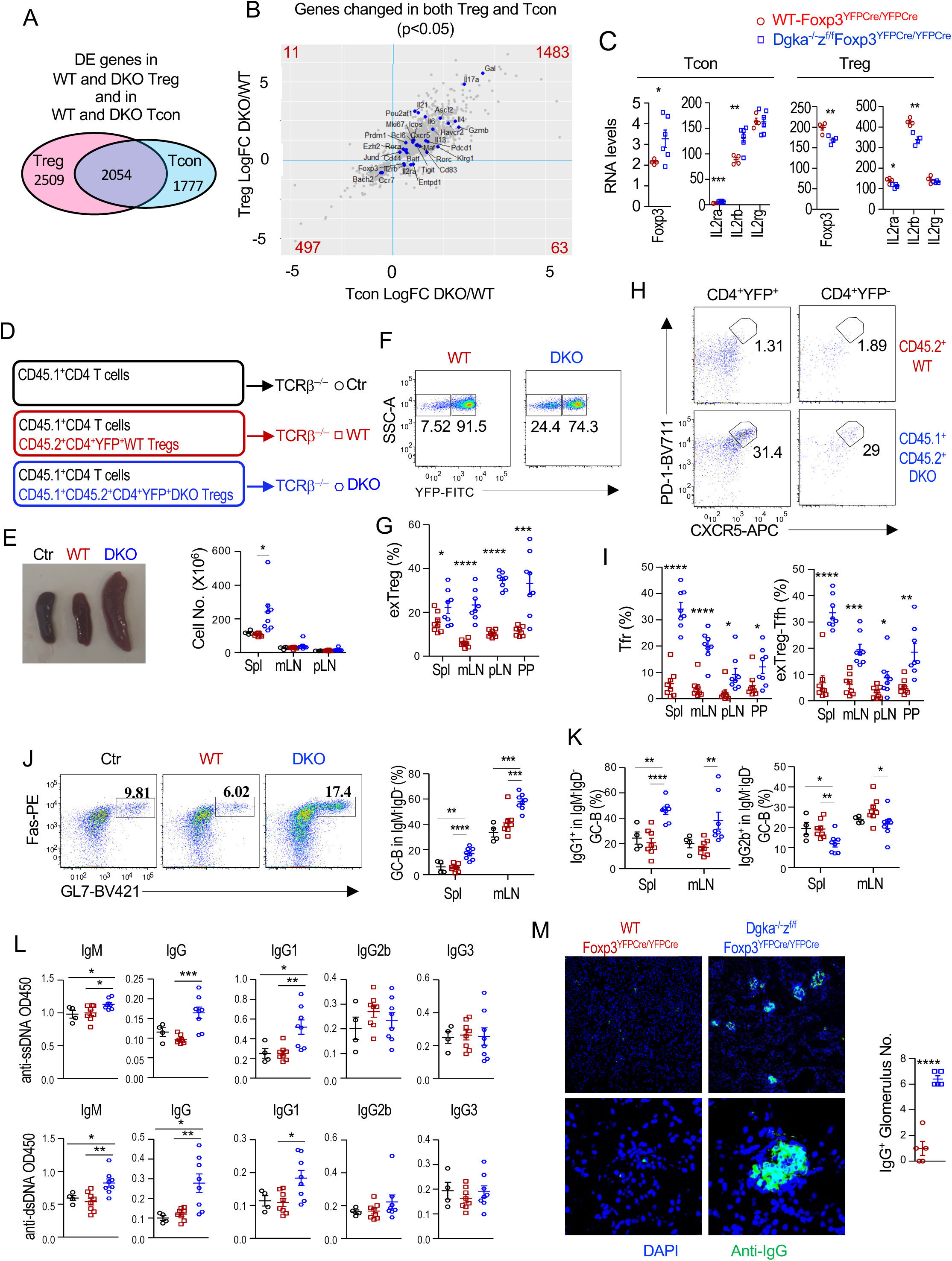
Enhanced conversion to exTreg/exTreg-Tfh cells and pathogenicity of αζDKO Tregs. **A–C.** Comparison of DEGs in both Treg-αζDKO Tregs and Tcons compared with their corresponding WT controls. **A.** DEGs that were shared or not shared in αζDKO Tregs and Tcons. **B.** Concordant and discordant expression of DEGs in Treg-αζDKO Tregs and Tcons. **C.** Expression of indicated Treg-associated genes in Treg-αζDKO Tregs and Tcons. **D - M.** Adoptive transfer experiments. 5 × 10^5^ double-sorted CD45.2^+^ Tregs from WT*-Foxp3^YFPCre/YFPCre^* or CD45.1^+^CD45.2^+^ Tregs from *DGKα^-/-^ζ^f/f^-Foxp3^YFPCre/YFPCre^*mice were co-injected *i.v.* with 1 × 10^6^ WT CD45.1^+^ CD4^+^ T cells into TCRβ^-/-^ mice. Serum were collected and mice were euthanized for the experiment 10 weeks after transfer. **D.** Experimental scheme. 5 × 10^5^ double-sorted CD45.2^+^ Tregs from WT*-Foxp3^YFPCre/YFPCre^* or CD45.1^+^CD45.2^+^ Tregs from *Dgka^-/-^z^f/f^-Foxp3^YFPCre/YFPCre^*mice were co-injected *i.v.* with 1 × 10^6^ WT CD45.1^+^ CD4^+^ T cells into TCRβ^-/-^ mice. Serum were collected and mice were euthanized for the experiment 10 weeks after transfer. **E.** Spleen sizes and total cell numbers (mean ± SEM). **F.** Representative FACS plot showing YFP levels in live gated CD45.2^+^ or CD45.1^+^CD45.2^+^CD4^+^TCRβ^+^ cells. **G.** Scatter plot showing means ± SEM of CD4^+^YFP^-^ exTreg percentages. **H.** Representative FACS plots showing PD-1 and CXCR5 staining in YFP^+^ 45.2^+^/CD45.1^+^CD45.2^+^CD4^+^TCRβ^+^ Tregs and YFP^-^ CD45.2^+^ or CD45.1^+^CD45.2^+^CD4^+^TCRβ^+^ exTregs. **I.** Mean ± SEM of Tfr and ExTreg-Tfh cell percentages. **J.** FACS plots showing Fas and GL7 staining in B220^+^ IgM^-^IgD^-^ B cells. Scatter plot showing mean ± SEM of GC-B cell percentages. **K.** IgG1^+^ and IgG2b^+^ cells in GC-B cells in the spleen and mLNs. **L.** Seral anti-ssDNA and -dsDNA antibody titers. **M.** IgG deposition in the kidney. Representative immunofluorescence of anti-IgG staining of kidney sections is shown. Scatter plot shows mean ± SEM of IgG^+^ glomerulus numbers per 10 × 10 field. Figure 10L is from one experiment and represents two experiments. Other data are representative of or pooled from 4–8 experiments. *, p < 0.05; **, p < 0.01; ***, p < 0.001; ****, p < 0.0001 by two-tailed unpaired Student t-test.

We initially used *Foxp3^YFPCre/YFPCre^-Rosa26^LSL-tdTomato^* reporter mice, an approach that has been used to map Treg fate, to further assess Treg to Tcon conversion. However, most *Foxp3^YFPCre/YFPCre^-Rosa26^LSL-tdTomato^* mice displayed variegated expression of tdTomato in many different cell types including CD8^+^ T-cells, B cells, CD4^+^Foxp3^-^ Tcons, and other immune cells in addition to Tregs among different mice (Supplemental Figure S13B). In extreme cases, the entire mouse could turn reddish, suggesting Cre-mediated germline recombination.

To further examine Treg-to-exTregs/exTreg-Tfh-cell conversion and to assess the pathogenicity of αζDKO Tregs/exTregs, we co-injected CD45.1^+^WT CD4^+^ T-cells (containing both Tregs and Tcons) with double-sorted CD4^+^YFP^+^TCRβ^+^ Tregs from either CD45.2^+^WT-*Foxp3^YFPCre/YFPCre^* or CD45.1^+^CD45.2^+^*Dgka^-/-^z^f/f^-Foxp3^YFPCre/YFPCre^* mice (both > 99% purity) into TCRβ^-/-^ mice (Figure 10D). Two to three months after transfer, recipients of αζDKO-Tregs manifested splenomegaly (Figure 10E), increased YFP^-^ exTregs within the transferred CD45.2^+^CD4^+^TCRβ^+^ population (Figure 10F, 10G), increased CXCR5^+^PD-1^+^ Tfr-cells within the YFP^+^ population, increased CXCR5^+^PD-1^+^ exTreg-Tfh-cells within the CD45.2^+^CD4^+^TCRβ^+^YFP^-^ exTregs (Figure 10H, 10I), and increased GC B-cells and IgG1^+^ cells within GC B-cells (Figure 10J, 10K). Moreover, αζDKO-Treg but not WT-Treg recipients developed elevated IgM- and IgG1-predominant autoantibodies (Figure 10L) and glomerular IgG deposition (Figure 10M). Increased αζDKO-Treg-to-exTreg conversion was also observed when both WT-Tregs and αζDKO-Tregs were co-injected into the same TCRβ^-/-^ hosts and thus in the same environment (Supplemental Figure S13C–S13F) or when they were injected individually into mildly irradiated WT hosts (Supplemental Figure S13G, S13H). Together, these data provided strong evidence that DGKαζ double deficiency accelerated Treg/Tfr-cell conversion to exTregs and exTreg-Tfh-cells. These data also suggested that DGKαζ deficient exTreg/exTreg-Tfh-cells may trigger deregulated GC B-cell and IgG1-predominant autoantibody responses. Of note, WT and αζDKO Tregs contained different ratios of effector Treg lineages and Tfr cells, which could partially contribute to the phenotypes observed. Future adoptive transfer experiments with individual Treg sublineages should provide additional insight into the mechanistic control of Treg to exTreg/Tfh cell conversion.

## Discussion

We have demonstrated that DGKαζ is critical for Tregs to maintain normal homeostasis and phenotypic and functional properties. In the absence of DGKαζ, Treg/Tcon ratios in lymphoid organs are reset to high levels due to enhanced Treg homeostatic proliferation; cTreg differentiation to eTreg is enhanced; and both cTregs and/or eTregs express abnormal levels of signature molecules. Increased CTLA-4, TIGIT, and ICOS may contribute to enhanced in vitro contact inhibition; increased PD-1 may reflect enhanced Treg activation and entry of functional exhaustion; decreased CD25 may destabilize Tregs (66–69). The abnormalities in Tregs and the development of multiorgan autoimmune diseases and increased effector CD8^+^ T-cells in Treg-αζDKO mice but not in DGKα or ζ single knockout mice suggest that DGKα and ζ synergistically prevent Tregs from functional impairment in vivo. However, due to the lack of a Treg specific DGKα deficient mouse model and the fact that Treg-αζDKO mice are DGKα germline deficiency, we could not completely rule out that DGKα deficiency in Tcons, CD8^+^ T cells, B cells, and/or other cells might contribute to the autoimmune phenotypes in the context of DGKαζ double deficiency in Tregs.

Tregs differentiate to multiple effector sublineages such as Treg1, Treg2, Treg17, and Tfr sublineages. However, proinflammatory cytokines and functions associated with Th-cells are usually suppressed in WT-Treg sublineages. In Treg-αζDKO mice, Treg2, Treg17, and Tfr-cells but not Treg1 cells are increased in numbers. At present, we do not know if the increases of Treg effector cells would enhance their pertinent suppressive functions. However, αζDKO Tregs express elevated Th-associated cytokines such as IFNγ, IL-17, and IL-4, suggesting that DGKαζ serves as a signal checkpoint to prevent Treg effector lineages differentiation and may also inhibit expression of cytokines that are usually associated with Th cells in Tregs. While we suspect that elevated proinflammatory cytokine expression by αζDKOTregs/exTregs may contribute to autoimmunity in Treg-αζDKO mice, further studies are needed to make firm conclusion.

Although Treg instability can lead to autoimmune diseases because of their self-reactivity (12, 13, 70), WT Treg-to-exTreg conversion is normally very limited in the steady state (10, 11, 42, 71, 72). Treg-to-exTreg-Tfh cell conversion has been rarely reported (73). A recent study identified that a CD25^low^ Tfr-cell population tends to lose function and become “ex-Tfr” cells (74). Mechanisms that ensure Treg stability are still not fully understood. Regulation of exTreg-Tfh-cell generation, functions, and pathogenicity is virtually unknown. We have revealed accelerated conversion of αζDKO-Tregs to exTregs, especially exTreg-Tfh-cells, and expansion of these cells in a CD28-independent manner. Upregulated ICOS may equip αζDKO exTreg-Tfh-cells with enhanced capability to receive positive-feedback signals from already increased GC B-cells to further promote expansion and function of exTreg-Tfh-cells. These αζDKO Tregs/exTreg cells are capable of triggering IgG1-predominant autoimmunity and GC B-cells. Our data indicate that normal Treg stability requires synergistical function of DGKα and ζ as DGKαζ activity serves as a signal checkpoint by braking Treg-to-exTreg/exTreg-Tfh cell conversion and by limiting numbers and functions of pathogenic IL4/13/21-expressing Tregs and exTreg/exTreg-Tfh cells to prevent IgG1-/IgE-predominant GC B-cell responses and/or autoimmunity.

DGKαζ double deficiency could affect Treg stability via multiple mechanisms. In αζDKO-Tregs, mammalian sterile 20-like kinase 1 (Mst1) (75), TET1/2/3 (76), and ring finger protein 31 (RNF31) (77), which promote Treg stability, were decreased; Pim-2 Kinase (78), cyclin-dependent kinase 2 (CDK2) (79), and tumor progression locus 2 (Tpl2) (80), which decrease Foxp3 and Treg stability, are increased (Supplemental Table 1). Foxo1 is important for Treg homeostasis and stability but inhibits Tfh- and Th17-cell differentiation (81); reduced Foxo activity may destabilize αζDKO-Tregs and promote their exTreg differentiation. CD25 is important for Treg stability and is downregulated in Tfr-cells, which appear less stable than other Tregs (68, 74). IL-2 signal also inhibits Tfh cell differentiation (82). Decreased CD25 in αζDKO-Tregs may destabilize them and accelerate their conversion to exTreg cells. Additionally, αζDKO-Tregs have altered metabolism such as enhanced glycolysis, are hyperproliferative, and upregulate many Th-/Tfh-associated TFs. Whether these abnormalities contribute to Treg destabilization remains to be investigated.

Signals from the TCR and CD28 are important for Treg maintenance and function (21, 22, 65). How these signals are regulated is not fully understood. We have revealed that DGKα and ζ impose Treg’s reliance on CD28 costimulatory signal. Treg homeostasis, Tfh-/Tfr-cell differentiation and function, and GC responses depend on CD28 (63–65). CD28 deficiency causes severely decreased Tregs; virtual absence of Tfh, Tfr, and GC B-cells; and decreased serum IgG1 but increased IgG3 levels. αζDKO in Tregs/exTregs partially or completely reverses these phenotypes caused by CD28 deficiency. Using both transcriptomic analysis and intracellular staining, we showed that DGKαζ controls multiple DAG-mediated pathways downstream of the TCR to exert their roles in Tregs. Transcriptomic analyses reveal that αζDKO Tregs display altered signaling and metabolic and transcription programs such as TCR signaling, MAPK, NFκB, Foxo, cell cycle, chemokine, and many other pathways. Intracellular staining further confirms enhanced Erk1/2 and mTORC1/2 activation in αζDKO Tregs. Many of these pathways are important Treg development, homeostasis, function, and/or stability. Enhanced DAG-Ras-Erk1/2 and mTOR signaling and enhanced uptake of glucose and glycolysis may trigger and fuel αζDKO-Treg proliferation, gain of proinflammatory function, and destabilization. Because both mTORC1 and mTORC2 are crucial for Tfh differentiation (83, 84), enhanced mTOR activity in αζDKO-Tregs/exTregs could also intrinsically promote exTreg-Tfh differentiation/expansion, which might further promote Tfh differentiation via bystander activating mechanism as observed in Tsc1 deficient mice (85). Future studies may illustrate how deregulation of each of these pathways may influence Treg stability and generation and function of exTreg-Tfh-cells as well as other aspects of Treg properties and functions.

In summary, we have demonstrated that DGKα and ζ play synergistic and critical roles in Tregs via tight control of multiple signaling pathways and metabolic and transcriptional programs. DGKα and ζ set normal Treg pool size by limiting their proliferation, restrain cTreg- to-eTreg differentiation, suppress Tregs’ proinflammatory programs, enforce Tregs’ dependence on CD28 costimulatory signal, and promote Tregs’ stability to brake Treg/Tfr cell-to-exTreg/exTreg-Tfh-cells conversion and subsequent expansion to prevent deregulated GC and autoantibody, especially IgG1/IgE predominant, responses. Our data provide the first evidence to suggest that exTreg-Tfh cells, at least in the Treg-αζDKO mouse model, are an important or even major source of pathogenic Tfh-cells that trigger deregulated autoantibody responses and autoimmune diseases. In line with many autoimmune diseases associated with deregulated Tfh and GC B-cells and elevated class-switched high affinity autoantibodies, future studies should determine whether exTreg-Tfh-cells play important roles in other autoimmune diseases in both animal models and human patients.

## Methods

### Experimental animals

C57BL/6, B6-CD45.1^+^, B6-Thy1.1^+^, *Foxp3^YFPCre^*, *Rosa26^LSL-tdTomato^*, *Tcrb^-/-^* mice, *Rag1^-/-^*, and *Cd28^-/-^* mice were purchased from the Jackson Laboratory. *Dgka^-/-^* and *Dgkz^f/f^*mice were previously reported (29, 31). Mice were maintained in specific pathogen-free facilities at Duke University. Both male and female mice were used for experiments. Single cells from the indicated organs were resuspended in IMDM supplemented with 10% FBS, 1% penicillin/streptomycin, and 50 mM 2-mercaptoethanol (IMDM-10) according to standard protocols.

### Antibodies, reagents, and flow cytometry

Antibodies and reagents are listed in Supplemental Table 6. Cells were stained with fluorescently conjugated antibodies in 2% FBS-PBS (PBS-2). Cell surface markers were stained at 4°C for 30 min. For intracellular staining of TFs, signaling molecules such as phospho-S6, phospho-Akt Ser473, and phospho-Erk, and other molecules such as YFP, Ki67, and CTLA-4, cells were fixed and permeabilized utilizing the Foxp3/TF Staining buffer set. Intracellular staining for IgG1 and IgG2b was performed by using the BD Biosciences Cytofix/Cytoperm and Perm/Wash solutions. For cell death, Live/Dead Fixable Violet Dead Cell Stain (Invitrogen, Carlsbad, CA) was used according to the manufacturer’s protocol. Stained cells were acquired on a FACS Fortessa or Canto II (BD Biosciences) device. FACS data were analyzed with FlowJo software (Version 9.9.6).

### CD4 T-cell enrichment and sorting

To enrich CD4^+^ T-cells, total splenocytes and LN cells were resuspended in 500 μl of IMDM-10 and incubated with 50 μl of CD4 MicroBeads (Miltenyi Biotec, Cat. No. 130-117-043) at 4°C for 30 minutes. After washed with IMDM-10, cells were resuspended in 2,000 μl PBS-2 and 1 mM EDTA. CD4^+^ T-cells were enriched using the LS columns (Miltenyi Biotec, Cat. No. 130-042-401). Enriched CD4^+^ cells were further stained with APC-anti-CD4, PE-Cy7-anti-CD8. Live CD4^+^CD8^-^YFP^+^ Tregs and CD4^+^CD8^-^YFP^-^ Tcons were sorted on an Aria II cell sorter (70-μm nozzle). Sorted cells were used to make RNA and DNA or double sorted for injection.

### In vitro stimulation

Splenocytes and LN cells were stimulated with 50 ng/ml PMA and 500 ng/ml ionomycin in the presence of brefedin A (1 ng/ml) for 4–5 hours. After cell surface staining, intracellular staining for Foxp3, GFP, IL17A, IL4, and IFNγ was performed using the EBiosciences Foxp3 Cytofix/Cytoperm and Perm/Wash solutions. For Treg proliferation, 2 × 10^5^ CTV-labeled splenocytes were seeded in a U-bottom 96-well plate and stimulated with an anti-CD3 antibody (145-2C11, 0.1 μg/ml) at 37°C for 72 hours.

### Treg-mediated contact inhibition

CTV-labeled CD45.1^+^CD4^+^CD25^-^ WT Tcons (5 × 10^4^) were mixed with CD45.2^+^CD8^-^ CD4^+^Foxp3YFP^+^ Tregs sorted from WT- or αζDKO-Foxp3^YFPCre/YFPCre^ mice at the ratios of 2:1 and 16:1 as well as 2 × 10^5^ mitomycin C-treated *Tcrb^-/-^*splenocytes and stimulated with an anti-CD3ε antibody (1 μg/ml). Cells were counted and stained for FACS analyses 72 hours later.

### BrdU incorporation assay

Mice were intraperitoneally injected with 1.5 mg of BrdU (150 μl of 10 mg/ml stock solution, company and cat no). Splenocytes and LN cells were stained for surface markers 8–10 hours after BrdU injection and then intracellular staining for BrdU according to the manufacturer’s protocol (BD Biosciences).

### Mixed BM chimeric mice

CD45.1^+^CD45.2^+^ WT mice were lethally irradiated (1,000 rad) and intravenously injected with a mixture of BM cells (1.5–2 X10^7^) from CD45.1^+^ WT and CD45.2^+^ WT*-Foxp3^YFPCre/YFPCre^*, *Dgka^-/-^z^f/f^*_-_*Foxp3^YFPCre/YFPCre^*, *Dgka^-/-^z^f/f^-Foxp3^YFPCre/YFPCre^-Cd28^-/-^*, or *Cd28^-/^*-*Foxp3^YFPCre/YFPCre^*mice. Chimeric mice were examined 8 weeks after irradiation.

### T cell adoptive transfer

CD45.2^+^ double-sorted WT or αζDKO Tregs (0.25 × 10^6^, purity > 99%) were *i.p.* injected with 0.5 × 10^6^ enriched CD45.1^+^ WT CD4^+^ T cells into TCRβ^-/-^ mice and analyzed for 10–12 weeks after adoptive transfer (Figure 10D–10M). For experiments in Supplemental Figure S11C–S11F, a mixture of double-sorted 0.25 × 10^6^ CD45.2^+^ WT Tregs and CD45.1^+^CD45.2^+^ αζDKO Tregs as well as 0.5 × 10^6^ CD45.1^+^ CD4 T cells were co-injected into TCRβ^-/-^ mice. Recipient mice were analyzed 8 weeks later. For experiments in Supplemental Figure S11G and S11H, 0.5 × 10^6^ CD45.2^+^ WT or αζDKO Tregs were injected into CD45.1^+^ WT mice 4 hours after mild irradiation (400 rad). Recipient mice were analyzed 14 days later.

### Glucose uptake assay

Two million splenocytes in 200 μl PBS from WT*-Foxp3^YFPCre/YFPCre^* and *Dgka^-/-^z^f/f^-Foxp3^YFPCre/YFPCre^* mice were seeded into a U-bottom 96-well plate in the presence or absence of 100 μM 2-(N-(7-Nitrobenz-2-oxa-1, 3-diazol-4-yl) Amino)-2-Deoxyglucose (2-NBDG; Life Technologies). After incubation at 37°C with 5% CO_2_ for 30min, 2-NBDG uptake was stopped by removing culture medium and washed with pre-chilled PBS twice. Cells were stained for surface markers before analysis with flow cytometry.

### Metabolic profiling

Sorted Tregs were washed with IMDM-10 and seeded in a 96-well plate at 2.5 × 10^5^ cells/well. Extracellular acidification rate (ECAR) and OCR were measured using an XFp Extracellular Flux Analyzer under glycolysis, mitochondrial stress, and mitochondrial fuel test conditions (Seahorse Bioscience/Agilent). For the glycolysis stress test, the assay buffer was made of nonbuffered DMEM medium supplemented with 2mM glutamine, and d-glucose, oligomycin, and 2DG were sequentially injected at final concentrations of 10mM, 1µM, and 50mM, respectively. For the mitochondrial stress test, the assay buffer was made of nonbuffered DMEM medium supplemented with 2.5mM d-glucose, 2mM glutamine, and 1mM sodium pyruvate, and oligomycin, carbonyl cyanide 4-(trifluoromethoxy)phenylhydrazone, and rotenone/antimycin A were sequentially injected at final concentrations of 1µM, 1µM, and 500nM, respectively. For the mitochondrial fuel test, the assay buffer was made of nonbuffered DMEM medium supplemented with 2.5mM d-glucose, 2mM glutamine, and 1mM sodium pyruvate, and UK5099, etomoxir, and bis-2-(5-phenylacetamido-1,3,4-thiadiazol-2-yl)ethyl sulfide were sequentially injected at final concentrations of 2µM, 4µM, and 3µM, respectively. Baseline ECAR (for the glycolysis stress test) and OCR (for the mitochondrial stress and mitochondrial fuel tests) values were averaged between technical replicates for these first three successive time intervals.

### Histology and IgG deposition in the kidney

Organs harvested from 6–16-months-old mice were fixed in 10% formalin overnight, preserved in 70% ethanol, and embedded in paraffin. The paraffin thin sections were stained with hematoxylin and eosin (H&E). Cryosections of kidneys frozen in OCT embedding medium were fixed with cold acetone for 5 min before rehydration in Tris-buffered saline (TBS, 50 mM Tris-Cl, pH 7.5, 150 mM NaCl). After being blocked with TBS containing 2% BSA and 5% normal donkey serum for 30 min, cryosections were incubated with an Alexa Fluor 488-anti-mouse IgG antibody at 4°C overnight. Slides were washed for 15 min in TBS before mounting in Vectashield hard set with DAPI.

### Enzyme-Linked Immunosorbent Assay and HEp-2 anti-nuclear antibody detection

Fifty microliters of appropriately diluted serum samples were added to 96-well plates precoated with anti-mouse Igκ and Igλ antibodies (2 mg/ml; Southern Biotech, Birmingham, AL) in 0.1 M carbonate buffer (pH 9.0). After incubation at 4°C overnight and multiple washes, total and subtype Ig concentrations were detected using HRP-conjugated goat anti-mouse anti-IgM, IgG, IgG1, IgG2b, IgG3, and IgE antibodies (Southern Biotech) with a TMB solution (Biolegend). For anti-dsDNA and anti-ssDNA antibodies, plates precoated with dsDNA or ssDNA were added with 1:30 diluted serum samples and were similarly detected with HRP-conjugated secondary antibodies.

To detect antinuclear antibodies, HEp-2 cells adhered to slides were fixed and permeabilized and incubated with 1:40 diluted serum samples. Cells were stained with goat anti-mouse IgG (H + L)–FITC (1:5000 dilution, Southern Biotech) and DAPI (500ng/ml, Thermo Fisher Scientific). Slides were imaged on a Leica SP5 confocal microscope.

### RNA-sequencing and transcriptomic analysis

Total RNA was isolated from CD4^+^Foxp3YFP^+^ Tregs and CD4^+^Foxp3YFP^-^ Tcons from WT*-Foxp3^YFPCre/YFPCre^* and *Dgka^-/-^z^fl/fl^-Foxp3^YFPCre/YFPCre^*mice (10–12-week-old) using the RNeasy Plus Mini kit (Qiagen). RNA-sequencing libraries were prepared using Truseq stranded mRNA kit (Illumina) according to the manufacturer’s protocol and sequenced at the Sequencing and Genomic Technologies Shared Resource at Duke University using the Illumina HiSeq2500 single end 50 bp platform. Raw sequencing reads were processed with FASTQC to screen eligible samples, and then single-end reads were trimmed for adaptor sequences and filtered with the Soapnuke tool. Trimmed reads were aligned by STAR (2.5.1b) to the mouse Ensembl genome (Ensembl, GRCm38.p5) with Ensembl annotation (Mus_musculus.GRCm38.84.gtf). Counting of reads on annotated transcripts was performed with htseq-count (0.6.0). The limma (3.38.3) Biconductor library has been used for counts normalization and differential analysis between the transcriptomes of the four WT and DKO Tregs. DEGs were subjected to enrichment analysis using ClusterProfiler (2.2.7). Biological processes in pathways in KEGG were chosen as significantly enriched terms with a p value less than 0.05.

### Statistical analysis

Statistical analyses were performed with Prism 5 (GraphPad). P values were calculated using two-tailed paired or unpaired Student’s t test. P values of less than 0.05 were considered significant. *p < 0.05, **p < 0.01, ***p < 0.001, and ****p < 0.0001 (Student’s t test).

### Study approval

All animal experiments were performed according to protocols approved by the IACUC of Duke University.

## Data availability

RNA sequencing data has been deposited in the GEO database (GSE276410). All original data associated with this manuscript are available from the contact author upon request.

## Supplemental material

The manuscript includes 13 supplemental figures and 6 supplemental tables with transcriptomic data.

## Author contributions

LL, HH, HXW, YP, TH, and SZ designed and performed experiments and analyzed data. PK and MBF performed data analysis and participated in manuscript preparation. LL prepared the figures and participated in manuscript preparation. JWS participated in data interpretation and manuscript preparation, and XPZ conceived the project, designed experiments, participated in data analysis, and wrote the paper.

## Acknowledgments

We thank the core facilities of Sequencing and Genomic Technologies, Flow Cytometry, Light Microscopy, and Cellular Metabolism Analysis, and the Research Immunohistology Lab at Duke University for their services and Jeffrey Zhong for editing the manuscript. Research in this manuscript is supported by the NIAID (R01AI079088, R56AI079088, and R01AI143781) and a Translating Duke Health Pilot Project Grant in Immunology.

## References

1. Sakaguchi S, Yamaguchi T, Nomura T, and Ono M. Regulatory T cells and immune tolerance. Cell. 2008;133(5):775–87.

2. Li MO, and Rudensky AY. T cell receptor signalling in the control of regulatory T cell differentiation and function. Nat Rev Immunol. 2016;16(4):220–33.

3. Bluestone JA, and Tang Q. Treg cells-the next frontier of cell therapy. Science. 2018;362(6411):154-5.

4. Hori S. Control of Regulatory T Cell Development by the Transcription Factor Foxp3. Science. 2003;299(5609):1057-61.

5. Fontenot JD, Gavin MA, and Rudensky AY. Foxp3 programs the development and function of CD4+CD25+ regulatory T cells. Nat Immunol. 2003;4(4):330–6.

6. Takahashi T, Kuniyasu Y, Toda M, Sakaguchi N, Itoh M, Iwata M, et al. Immunologic self-tolerance maintained by CD25+CD4+ naturally anergic and suppressive T cells: induction of autoimmune disease by breaking their anergic/suppressive state. Int Immunol. 1998;10(12):1969–80.

7. Samstein RM, Arvey A, Josefowicz SZ, Peng X, Reynolds A, Sandstrom R, et al. Foxp3 exploits a pre-existent enhancer landscape for regulatory T cell lineage specification. Cell. 2012;151(1):153–66.

8. Bettini ML, Pan F, Bettini M, Finkelstein D, Rehg JE, Floess S, et al. Loss of epigenetic modification driven by the Foxp3 transcription factor leads to regulatory T cell insufficiency. Immunity. 2012;36(5):717–30.

9. Zheng Y, Josefowicz S, Chaudhry A, Peng XP, Forbush K, and Rudensky AY. Role of conserved non-coding DNA elements in the Foxp3 gene in regulatory T-cell fate. Nature. 2010;463(7282):808-12.

10. Hori S. Lineage stability and phenotypic plasticity of Foxp3(+) regulatory T cells. Immunol Rev. 2014;259(1):159–72.

11. Sakaguchi S, Vignali DA, Rudensky AY, Niec RE, and Waldmann H. The plasticity and stability of regulatory T cells. Nat Rev Immunol. 2013;13(6):461–7.

12. Zhou X, Bailey-Bucktrout SL, Jeker LT, Penaranda C, Martinez-Llordella M, Ashby M, et al. Instability of the transcription factor Foxp3 leads to the generation of pathogenic memory T cells in vivo. Nat Immunol. 2009;10(9):1000–7.

13. Bailey-Bucktrout SL, Martinez-Llordella M, Zhou X, Anthony B, Rosenthal W, Luche H, et al. Self-antigen-driven activation induces instability of regulatory T cells during an inflammatory autoimmune response. Immunity. 2013;39(5):949–62.

14. DeFranco AL. Germinal centers and autoimmune disease in humans and mice. Immunol Cell Biol. 2016;94(10):918–24.

15. Crotty S. A brief history of T cell help to B cells. Nat Rev Immunol. 2015;15(3):185–9.

16. Arkatkar T, Du SW, Jacobs HM, Dam EM, Hou B, Buckner JH, et al. B cell-derived IL-6 initiates spontaneous germinal center formation during systemic autoimmunity. J Exp Med. 2017;214(11):3207–17.

17. Choi SC, Hutchinson TE, Titov AA, Seay HR, Li S, Brusko TM, et al. The Lupus Susceptibility Gene Pbx1 Regulates the Balance between Follicular Helper T Cell and Regulatory T Cell Differentiation. J Immunol. 2016;197(2):458–69.

18. Sage PT, and Sharpe AH. T follicular regulatory cells. Immunol Rev. 2016;271(1):246–59.

19. Fu W, Liu X, Lin X, Feng H, Sun L, Li S, et al. Deficiency in T follicular regulatory cells promotes autoimmunity. J Exp Med. 2018;215(3):815–25.

20. Jacquemin C, Schmitt N, Contin-Bordes C, Liu Y, Narayanan P, Seneschal J, et al. OX40 Ligand Contributes to Human Lupus Pathogenesis by Promoting T Follicular Helper Response. Immunity. 2015;42(6):1159–70.

21. Delpoux A, Yakonowsky P, Durand A, Charvet C, Valente M, Pommier A, et al. TCR signaling events are required for maintaining CD4 regulatory T cell numbers and suppressive capacities in the periphery. J Immunol. 2014;193(12):5914–23.

22. Levine AG, Arvey A, Jin W, and Rudensky AY. Continuous requirement for the TCR in regulatory T cell function. Nat Immunol. 2014;15(11):1070–8.

23. Gorentla BK, Wan CK, and Zhong XP. Negative regulation of mTOR activation by diacylglycerol kinases. Blood. 2011;117(15):4022–31.

24. Chen SS, Hu Z, and Zhong XP. Diacylglycerol Kinases in T Cell Tolerance and Effector Function. Front Cell Dev Biol. 2016;4:130.

25. Merida I, Arranz-Nicolas J, Rodriguez-Rodriguez C, and Avila-Flores A. Diacylglycerol kinase control of protein kinase C. Biochem J. 2019;476(8):1205–19.

26. Zhong XP, Hainey EA, Olenchock BA, Jordan MS, Maltzman JS, Nichols KE, et al. Enhanced T cell responses due to diacylglycerol kinase zeta deficiency. Nat Immunol. 2003;4(9):882–90.

27. Zha Y, Marks R, Ho AW, Peterson AC, Janardhan S, Brown I, et al. T cell anergy is reversed by active Ras and is regulated by diacylglycerol kinase-alpha. Nat Immunol. 2006;7(11):1166–73.

28. Guo R, Wan CK, Carpenter JH, Mousallem T, Boustany RM, Kuan CT, et al. Synergistic control of T cell development and tumor suppression by diacylglycerol kinase alpha and zeta. Proc Natl Acad Sci U S A. 2008;105(33):11909–14.

29. Olenchock BA, Guo R, Carpenter JH, Jordan M, Topham MK, Koretzky GA, et al. Disruption of diacylglycerol metabolism impairs the induction of T cell anergy. Nat Immunol. 2006;7(11):1174–81.

30. Baldanzi G, Pighini A, Bettio V, Rainero E, Traini S, Chianale F, et al. SAP-mediated inhibition of diacylglycerol kinase alpha regulates TCR-induced diacylglycerol signaling. J Immunol. 2011;187(11):5941–51.

31. Yang J, Zhang P, Krishna S, Wang J, Lin X, Huang H, et al. Unexpected positive control of NFkappaB and miR-155 by DGKalpha and zeta ensures effector and memory CD8+ T cell differentiation. Oncotarget. 2016;7(23):33744–64.

32. Ruffo E, Malacarne V, Larsen SE, Das R, Patrussi L, Wulfing C, et al. Inhibition of diacylglycerol kinase alpha restores restimulation-induced cell death and reduces immunopathology in XLP-1. Sci Transl Med. 2016;8(321):321ra7.

33. Jung IY, Kim YY, Yu HS, Lee M, Kim S, and Lee J. CRISPR/Cas9-Mediated Knockout of DGK Improves Antitumor Activities of Human T Cells. Cancer Res. 2018;78(16):4692–703.

34. Riese MJ, Wang LC, Moon EK, Joshi RP, Ranganathan A, June CH, et al. Enhanced effector responses in activated CD8+ T cells deficient in diacylglycerol kinases. Cancer Res. 2013;73(12):3566–77.

35. Shen S, Wu J, Srivatsan S, Gorentla BK, Shin J, Xu L, et al. Tight regulation of diacylglycerol-mediated signaling is critical for proper invariant NKT cell development. J Immunol. 2011;187(5):2122–9.

36. Pan Y, Deng W, Xie J, Zhang S, Wan ECK, Li L, et al. Graded diacylglycerol kinases alpha and zeta activities ensure mucosal-associated invariant T-cell development in mice. Eur J Immunol. 2020;50(2):192–204.

37. Yang J, Wang HX, Xie J, Li L, Wang J, Wan ECK, et al. DGK alpha and zeta Activities Control TH1 and TH17 Cell Differentiation. Front Immunol. 2019;10:3048.

38. Schmidt AM, Zou T, Joshi RP, Leichner TM, Pimentel MA, Sommers CL, et al. Diacylglycerol kinase zeta limits the generation of natural regulatory T cells. Sci Signal. 2013;6(303):ra101.

39. Joshi RP, Schmidt AM, Das J, Pytel D, Riese MJ, Lester M, et al. The zeta isoform of diacylglycerol kinase plays a predominant role in regulatory T cell development and TCR-mediated ras signaling. Sci Signal. 2013;6(303):ra102.

40. Bayer AL, Lee JY, de la Barrera A, Surh CD, and Malek TR. A function for IL-7R for CD4+CD25+Foxp3+ T regulatory cells. J Immunol. 2008;181(1):225–34.

41. DuPage M, Chopra G, Quiros J, Rosenthal WL, Morar MM, Holohan D, et al. The chromatin-modifying enzyme Ezh2 is critical for the maintenance of regulatory T cell identity after activation. Immunity. 2015;42(2):227–38.

42. Levine AG, Medoza A, Hemmers S, Moltedo B, Niec RE, Schizas M, et al. Stability and function of regulatory T cells expressing the transcription factor T-bet. Nature. 2017;546(7658):421–5.

43. Wang Y, Su MA, and Wan YY. An essential role of the transcription factor GATA-3 for the function of regulatory T cells. Immunity. 2011;35(3):337–48.

44. Zheng Y, Chaudhry A, Kas A, deRoos P, Kim JM, Chu TT, et al. Regulatory T-cell suppressor program co-opts transcription factor IRF4 to control T(H)2 responses. Nature. 2009;458(7236):351–6.

45. Chaudhry A, Rudra D, Treuting P, Samstein RM, Liang Y, Kas A, et al. CD4+ regulatory T cells control TH17 responses in a Stat3-dependent manner. Science. 2009;326(5955):986–91.

46. Chung Y, Tanaka S, Chu F, Nurieva RI, Martinez GJ, Rawal S, et al. Follicular regulatory T cells expressing Foxp3 and Bcl-6 suppress germinal center reactions. Nat Med. 2011;17(8):983–8.

47. Xu C, Fu Y, Liu S, Trittipo J, Lu X, Qi R, et al. BATF Regulates T Regulatory Cell Functional Specification and Fitness of Triglyceride Metabolism in Restraining Allergic Responses. J Immunol. 2021;206(9):2088–100.

48. Shan F, Cillo AR, Cardello C, Yuan DY, Kunning SR, Cui J, et al. Integrated BATF transcriptional network regulates suppressive intratumoral regulatory T cells. Sci Immunol. 2023;8(87):eadf6717.

49. Roychoudhuri R, Hirahara K, Mousavi K, Clever D, Klebanoff CA, Bonelli M, et al. BACH2 represses effector programs to stabilize T(reg)-mediated immune homeostasis. Nature. 2013;498(7455):506–10.

50. Hatzi K, Nance JP, Kroenke MA, Bothwell M, Haddad EK, Melnick A, et al. BCL6 orchestrates Tfh cell differentiation via multiple distinct mechanisms. J Exp Med. 2015;212(4):539–53.

51. Liu X, Chen X, Zhong B, Wang A, Wang X, Chu F, et al. Transcription factor achaete-scute homologue 2 initiates follicular T-helper-cell development. Nature. 2014;507(7493):513–8.

52. Stauss D, Brunner C, Berberich-Siebelt F, Hopken UE, Lipp M, and Muller G. The transcriptional coactivator Bob1 promotes the development of follicular T helper cells via Bcl6. EMBO J. 2016;35(8):881–98.

53. Gowthaman U, Chen JS, Zhang B, Flynn WF, Lu Y, Song W, et al. Identification of a T follicular helper cell subset that drives anaphylactic IgE. Science. 2019;365(6456).

54. Rao DA, Gurish MF, Marshall JL, Slowikowski K, Fonseka CY, Liu Y, et al. Pathologically expanded peripheral T helper cell subset drives B cells in rheumatoid arthritis. Nature. 2017;542(7639):110-4.

55. Yoshitomi H, and Ueno H. Shared and distinct roles of T peripheral helper and T follicular helper cells in human diseases. Cell Mol Immunol. 2021;18(3):523–7.

56. Bocharnikov AV, Keegan J, Wacleche VS, Cao Y, Fonseka CY, Wang G, et al. PD-1hiCXCR5-T peripheral helper cells promote B cell responses in lupus via MAF and IL-21. JCI Insight. 2019;4(20).

57. Ekman I, Ihantola EL, Viisanen T, Rao DA, Nanto-Salonen K, Knip M, et al. Circulating CXCR5(-)PD-1(hi) peripheral T helper cells are associated with progression to type 1 diabetes. Diabetologia. 2019;62(9):1681–8.

58. Selb R, Eckl-Dorna J, Neunkirchner A, Schmetterer K, Marth K, Gamper J, et al. CD23 surface density on B cells is associated with IgE levels and determines IgE-facilitated allergen uptake, as well as activation of allergen-specific T cells. J Allergy Clin Immunol. 2017;139(1):290–9 e4.

59. Ritvo PG, Churlaud G, Quiniou V, Florez L, Brimaud F, Fourcade G, et al. T(fr) cells lack IL-2Ralpha but express decoy IL-1R2 and IL-1Ra and suppress the IL-1-dependent activation of T(fh) cells. Sci Immunol. 2017;2(15).

60. Kumar S, Fonseca VR, Ribeiro F, Basto AP, Agua-Doce A, Monteiro M, et al. Developmental bifurcation of human T follicular regulatory cells. Sci Immunol. 2021;6(59).

61. Botta D, Fuller MJ, Marquez-Lago TT, Bachus H, Bradley JE, Weinmann AS, et al. Dynamic regulation of T follicular regulatory cell responses by interleukin 2 during influenza infection. Nat Immunol. 2017;18(11):1249–60.

62. Jacobsen JT, Hu W, TB RC, Solem S, Galante A, Lin Z, et al. Expression of Foxp3 by T follicular helper cells in end-stage germinal centers. Science. 2021;373(6552).

63. Walker LS, Gulbranson-Judge A, Flynn S, Brocker T, Raykundalia C, Goodall M, et al. Compromised OX40 function in CD28-deficient mice is linked with failure to develop CXC chemokine receptor 5-positive CD4 cells and germinal centers. J Exp Med. 1999;190(8):1115–22.

64. Tai X, Cowan M, Feigenbaum L, and Singer A. CD28 costimulation of developing thymocytes induces Foxp3 expression and regulatory T cell differentiation independently of interleukin 2. Nat Immunol. 2005;6(2):152–62.

65. Zhang R, Sage PT, Finn K, Huynh A, Blazar BR, Marangoni F, et al. B Cells Drive Autoimmunity in Mice with CD28-Deficient Regulatory T Cells. J Immunol. 2017;199(12):3972–80.

66. Chen Y, Shen S, Gorentla BK, Gao J, and Zhong XP. Murine regulatory T cells contain hyperproliferative and death-prone subsets with differential ICOS expression. J Immunol. 2012;188(4):1698–707.

67. Joller N, Lozano E, Burkett PR, Patel B, Xiao S, Zhu C, et al. Treg cells expressing the coinhibitory molecule TIGIT selectively inhibit proinflammatory Th1 and Th17 cell responses. Immunity. 2014;40(4):569–81.

68. Toomer KH, Lui JB, Altman NH, Ban Y, Chen X, and Malek TR. Essential and non-overlapping IL-2Ralpha-dependent processes for thymic development and peripheral homeostasis of regulatory T cells. Nat Commun. 2019;10(1):1037.

69. Tan CL, Kuchroo JR, Sage PT, Liang D, Francisco LM, Buck J, et al. PD-1 restraint of regulatory T cell suppressive activity is critical for immune tolerance. J Exp Med. 2021;218(1).

70. Komatsu N, Okamoto K, Sawa S, Nakashima T, Oh-hora M, Kodama T, et al. Pathogenic conversion of Foxp3+ T cells into TH17 cells in autoimmune arthritis. Nat Med. 2014;20(1):62–8.

71. Miyao T, Floess S, Setoguchi R, Luche H, Fehling HJ, Waldmann H, et al. Plasticity of Foxp3(+) T cells reflects promiscuous Foxp3 expression in conventional T cells but not reprogramming of regulatory T cells. Immunity. 2012;36(2):262–75.

72. Rubtsov YP, Niec RE, Josefowicz S, Li L, Darce J, Mathis D, et al. Stability of the regulatory T cell lineage in vivo. Science. 2010;329(5999):1667–71.

73. Tsuji M, Komatsu N, Kawamoto S, Suzuki K, Kanagawa O, Honjo T, et al. Preferential generation of follicular B helper T cells from Foxp3+ T cells in gut Peyer’s patches. Science. 2009;323(5920):1488-92.

74. Hou S, Clement RL, Diallo A, Blazar BR, Rudensky AY, Sharpe AH, et al. FoxP3 and Ezh2 regulate Tfr cell suppressive function and transcriptional program. J Exp Med. 2019;216(3):605–20.

75. Li J, Du X, Shi H, Deng K, Chi H, and Tao W. Mammalian Sterile 20-like Kinase 1 (Mst1) Enhances the Stability of Forkhead Box P3 (Foxp3) and the Function of Regulatory T Cells by Modulating Foxp3 Acetylation. J Biol Chem. 2015;290(52):30762–70.

76. Yue XJ, Trifari S, Aijo T, Tsagaratou A, Pastor WA, Zepeda-Martinez JA, et al. Control of Foxp3 stability through modulation of TET activity. Journal of Experimental Medicine. 2016;213(3):377–97.

77. Luo X, Nie J, Wang S, Chen Z, Chen W, Li D, et al. Poly(ADP-ribosyl)ation of FOXP3 protein mediated by PARP-1 regulates the function of regulatory T cells. J Biol Chem. 2016;291(3):1201.

78. Deng G, Nagai Y, Xiao Y, Li Z, Dai S, Ohtani T, et al. Pim-2 Kinase Influences Regulatory T Cell Function and Stability by Mediating Foxp3 Protein N-terminal Phosphorylation. J Biol Chem. 2015;290(33):20211–20.

79. Morawski PA, Mehra P, Chen C, Bhatti T, and Wells AD. Foxp3 protein stability is regulated by cyclin-dependent kinase 2. J Biol Chem. 2013;288(34):24494–502.

80. Li X, Acuff NV, Peeks AR, Kirkland R, Wyatt KD, Nagy T, et al. Tumor Progression Locus 2 (Tpl2) Activates the Mammalian Target of Rapamycin (mTOR) Pathway, Inhibits Forkhead Box P3 (FoxP3) Expression, and Limits Regulatory T Cell (Treg) Immunosuppressive Functions. Journal of Biological Chemistry. 2016;291(32):16802–15.

81. Kerdiles YM, Stone EL, Beisner DR, McGargill MA, Ch’en IL, Stockmann C, et al. Foxo transcription factors control regulatory T cell development and function. Immunity. 2010;33(6):890–904.

82. Ballesteros-Tato A, León B, Graf Beth A, Moquin A, Adams Pamela S, Lund Frances E, et al. Interleukin-2 Inhibits Germinal Center Formation by Limiting T Follicular Helper Cell Differentiation. Immunity. 2012;36(5):847–56.

83. Yang J, Lin X, Pan Y, Wang J, Chen P, Huang H, et al. Critical roles of mTOR Complex 1 and 2 for T follicular helper cell differentiation and germinal center responses. Elife. 2016;5.

84. Zeng H, Cohen S, Guy C, Shrestha S, Neale G, Brown SA, et al. mTORC1 and mTORC2 Kinase Signaling and Glucose Metabolism Drive Follicular Helper T Cell Differentiation. Immunity. 2016;45(3):540–54.

85. Zhang S, Li L, Xie D, Reddy S, Sleasman JW, Ma L, et al. Regulation of Intrinsic and Bystander T Follicular Helper Cell Differentiation and Autoimmunity by Tsc1. Front Immunol. 2021;12:620437.

