## Supplemental figure S1 for "DGKα and ζ Deficiency Causes Regulatory T-Cell Dysregulation, Destabilization, and Conversion to Pathogenic T-Follicular Helper Cells to Trigger IgG1-Predominant Autoimmunity"

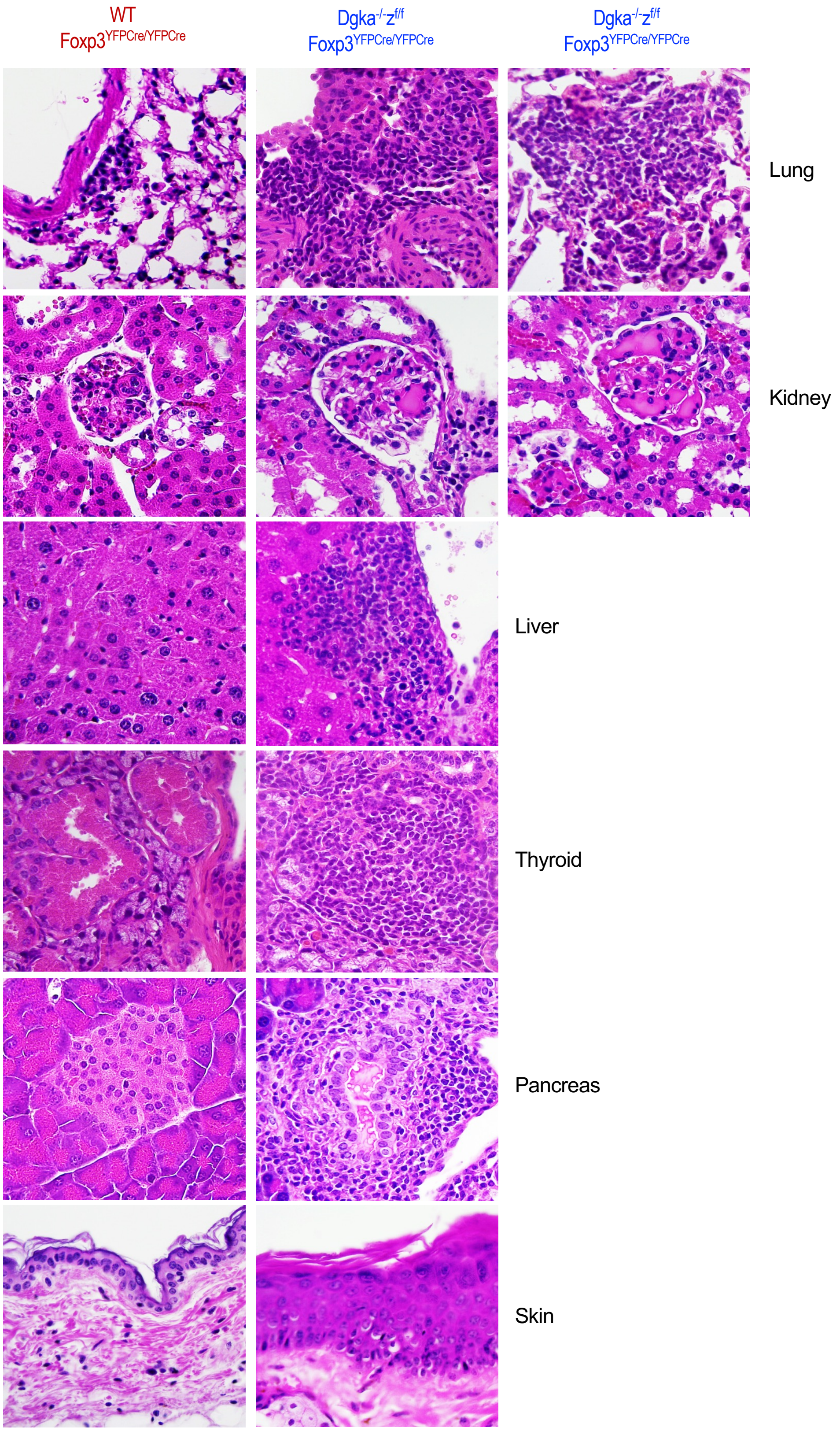


**Supplemental Figure S1. H&E staining of indicated organs correlating to Figure 1D with high magnification of the rectangle areas.**
