## Supplemental figure S2 for "DGKα and ζ Deficiency Causes Regulatory T-Cell Dysregulation, Destabilization, and Conversion to Pathogenic T-Follicular Helper Cells to Trigger IgG1-Predominant Autoimmunity"

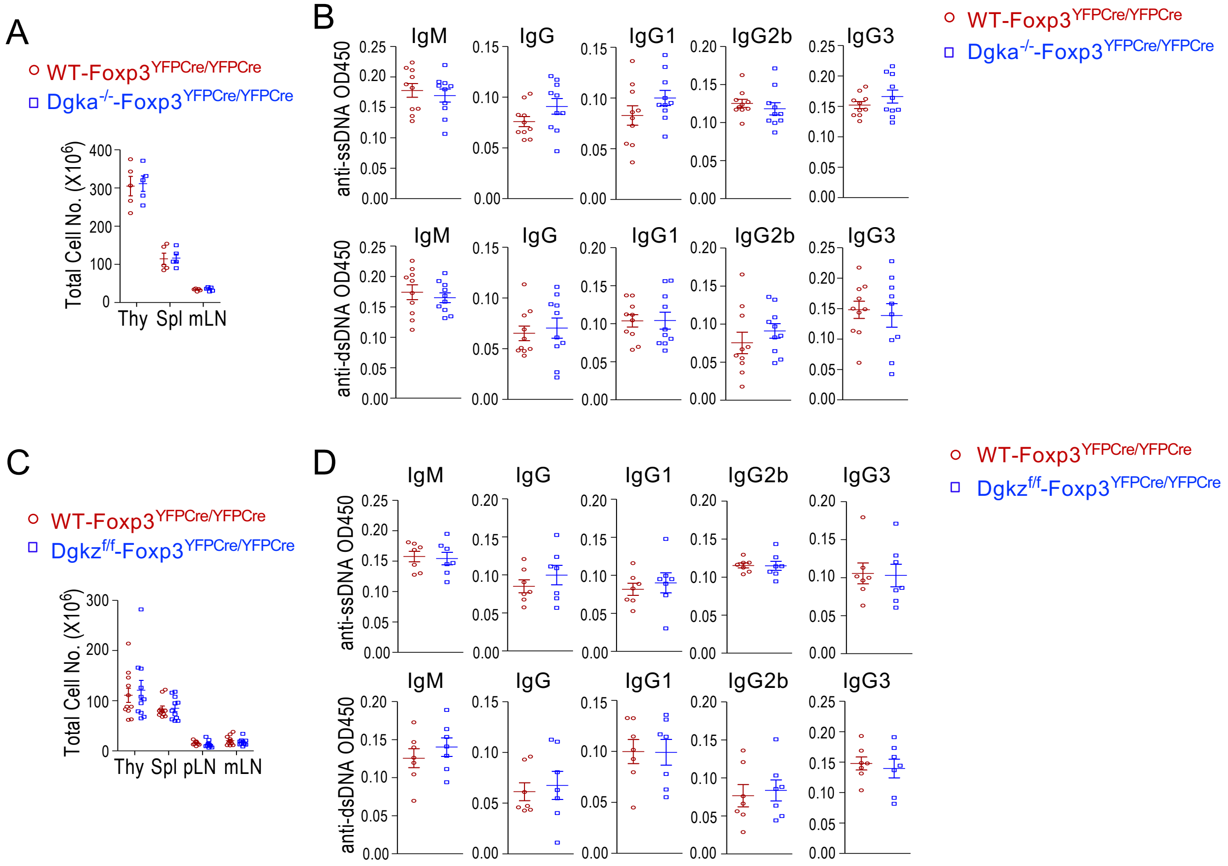


**Supplemental Figure S2. Analyses of *$Dgka^{-/-}$-$Foxp3^{YFPCre/YFPCre}$* mice and *$Dgka^{+/+}z^{f/f}$-$Foxp3^{YFPCre/YFPCre}$* mice. A–B.** *$Dgka^{-/-}$-$Foxp3^{YFPCre/YFPCre}$* mice and WT-*$Foxp3^{YFPCre/YFPCre}$* mice. **A.** Total cellularity. **B.** Serum autoantibodies. **C–D.** *$Dgka^{+/+}z^{f/f}$-$Foxp3^{YFPCre/YFPCre}$* mice and WT-*$Foxp3^{YFPCre/YFPCre}$* mice. **C.** Total cellularity. **D.** Serum autoantibodies. Each circle or square represents one mouse of the indicated genotypes. Data shown are presentative of or pooled from at least five experiments.
