## Supplemental figure S3 for "DGKα and ζ Deficiency Causes Regulatory T-Cell Dysregulation, Destabilization, and Conversion to Pathogenic T-Follicular Helper Cells to Trigger IgG1-Predominant Autoimmunity"

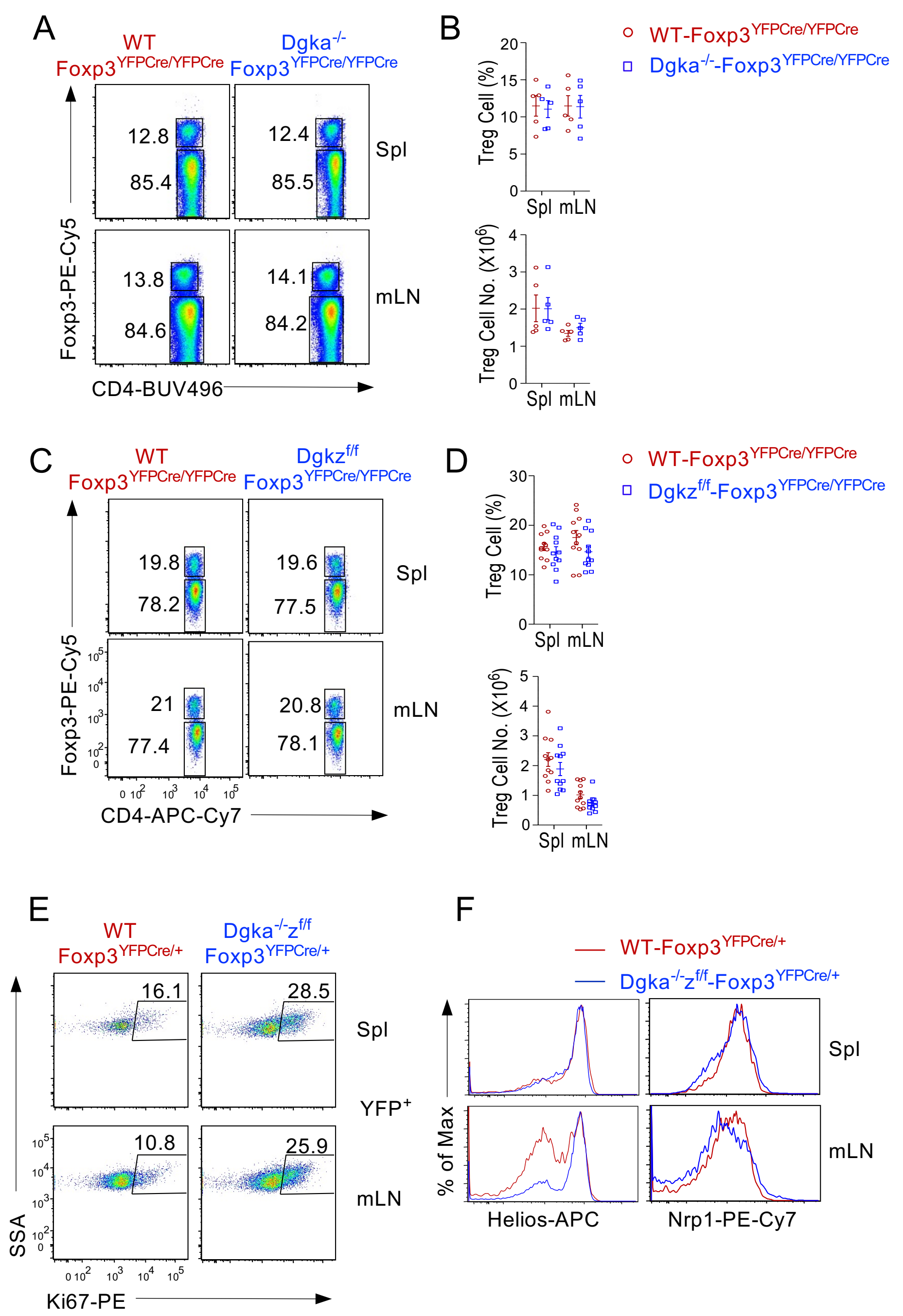


**Supplemental Figure S3. Analyses of Tregs in DGKα or DGKζ single knockout mice. A, B.** Tregs in Dgka$^{-/-}$ mice. **C, D.** Tregs of *Dgka$^{+/+}$z$^{f/f}$-Foxp3$^{YFPCre/YFPCre}$* mice. **E, F.** Analysis of female *Dgka$^{-/-}$z$^{f/f}$-Foxp3$^{YFPCre/+}$* and WT-*Foxp3$^{YFPCre/+}$* mice. **E.** Ki67 staining in YFP$^{+}$ Tregs. **F.** Overlaid histograms showing Helios and Nrp1 expression in YFP$^{+}$ Tregs. Each circle or square represents one mouse of the indicated genotypes. Data shown are presentative of or pooled from at least five experiments.
