## Supplemental figure S4 for "DGKα and ζ Deficiency Causes Regulatory T-Cell Dysregulation, Destabilization, and Conversion to Pathogenic T-Follicular Helper Cells to Trigger IgG1-Predominant Autoimmunity"

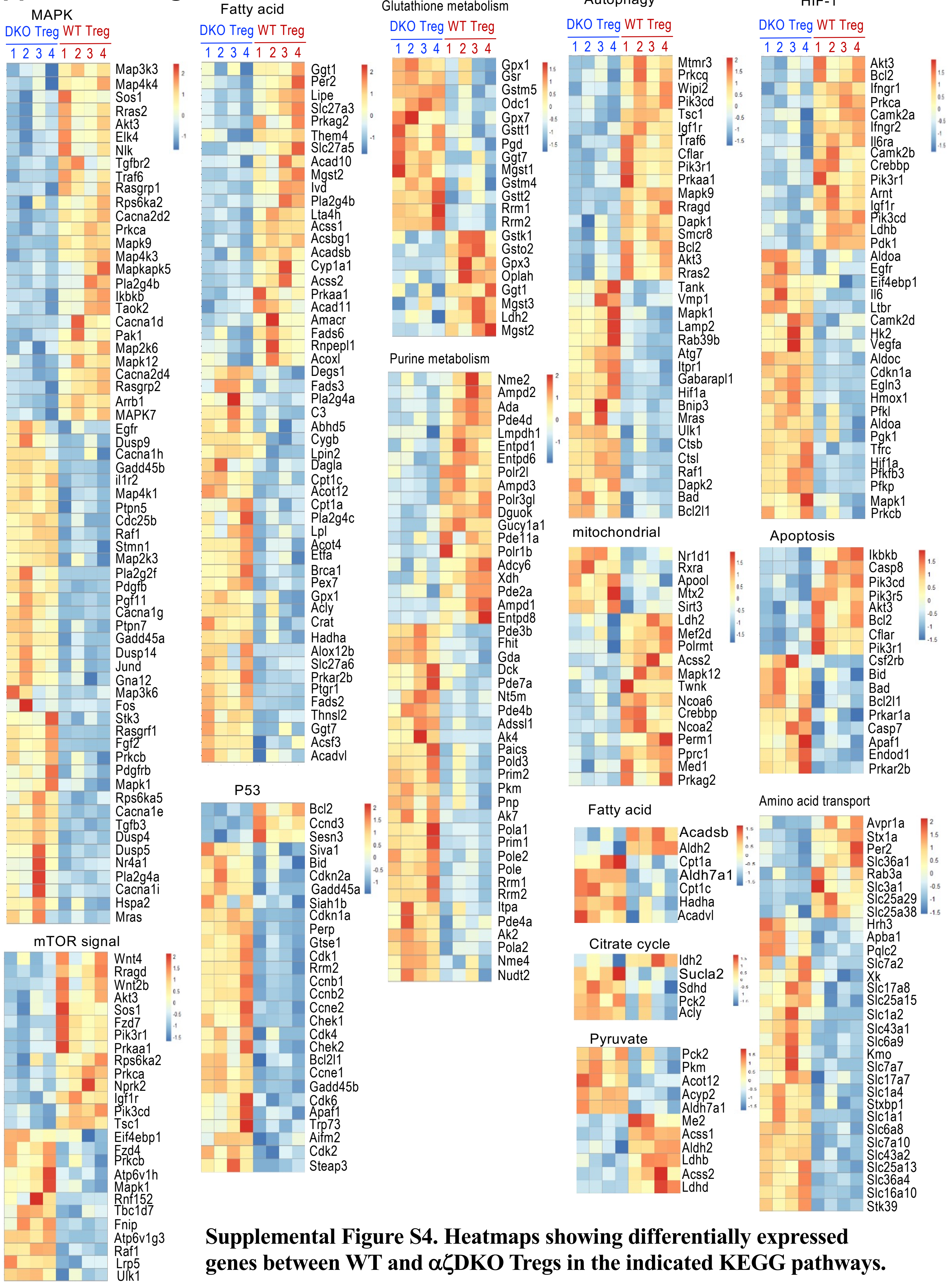


**Supplemental Figure S4. Heatmaps showing differentially expressed genes between WT and αζDKO Tregs in the indicated KEGG pathways.**
