## Supplemental figure S5 for "DGKα and ζ Deficiency Causes Regulatory T-Cell Dysregulation, Destabilization, and Conversion to Pathogenic T-Follicular Helper Cells to Trigger IgG1-Predominant Autoimmunity"

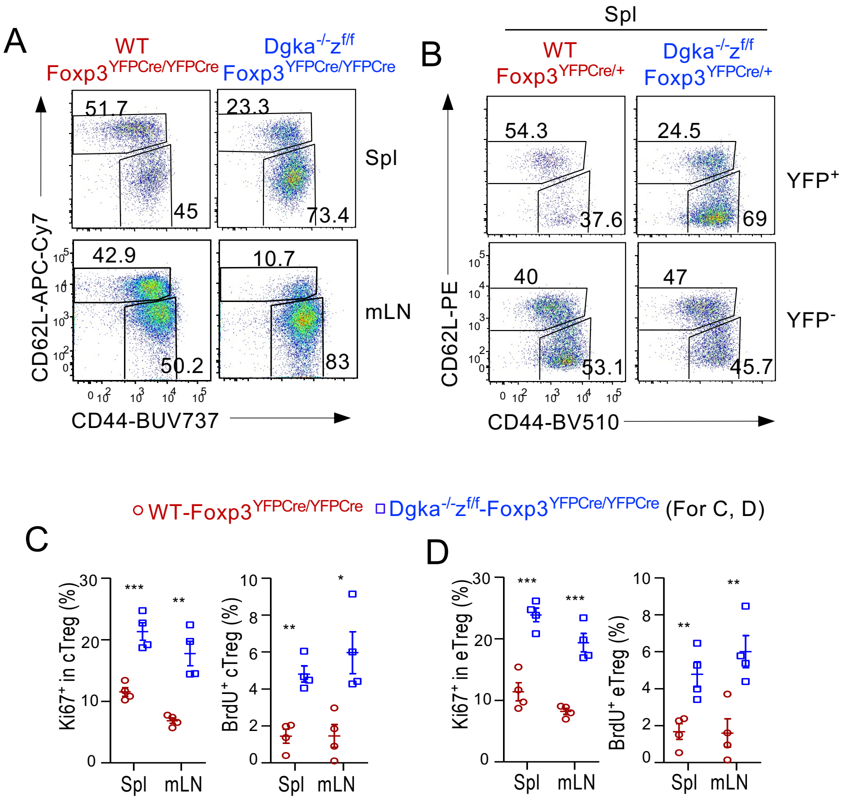


**Supplemental Figure S5. A.** CD44 and CD62L expression in CD4$^+$Foxp3$^+$ Tregs from *Dgka$^{-/-}$z$^{f/f}$-Foxp3$^{YFPCre/YFPCre}$* and WT-*Foxp3$^{YFPCre/YFPCre}$* control mice. **B.** CD44 and CD62L expression in YFP$^+$ and YFP$^-$ CD4$^+$Foxp3$^+$ Tregs from female *Dgka$^{-/-}$z$^{f/f}$-Foxp3$^{YFPCre/+}$* and WT- *Foxp3$^{YFPCre/+}$* control mice. **C, D.** Scatter plots show mean ± SEM of percentages of Ki67$^+$ or BrdU$^+$ cells within cTregs (C) and eTregs (D).
