## Supplemental figure S6 for "DGKα and ζ Deficiency Causes Regulatory T-Cell Dysregulation, Destabilization, and Conversion to Pathogenic T-Follicular Helper Cells to Trigger IgG1-Predominant Autoimmunity"

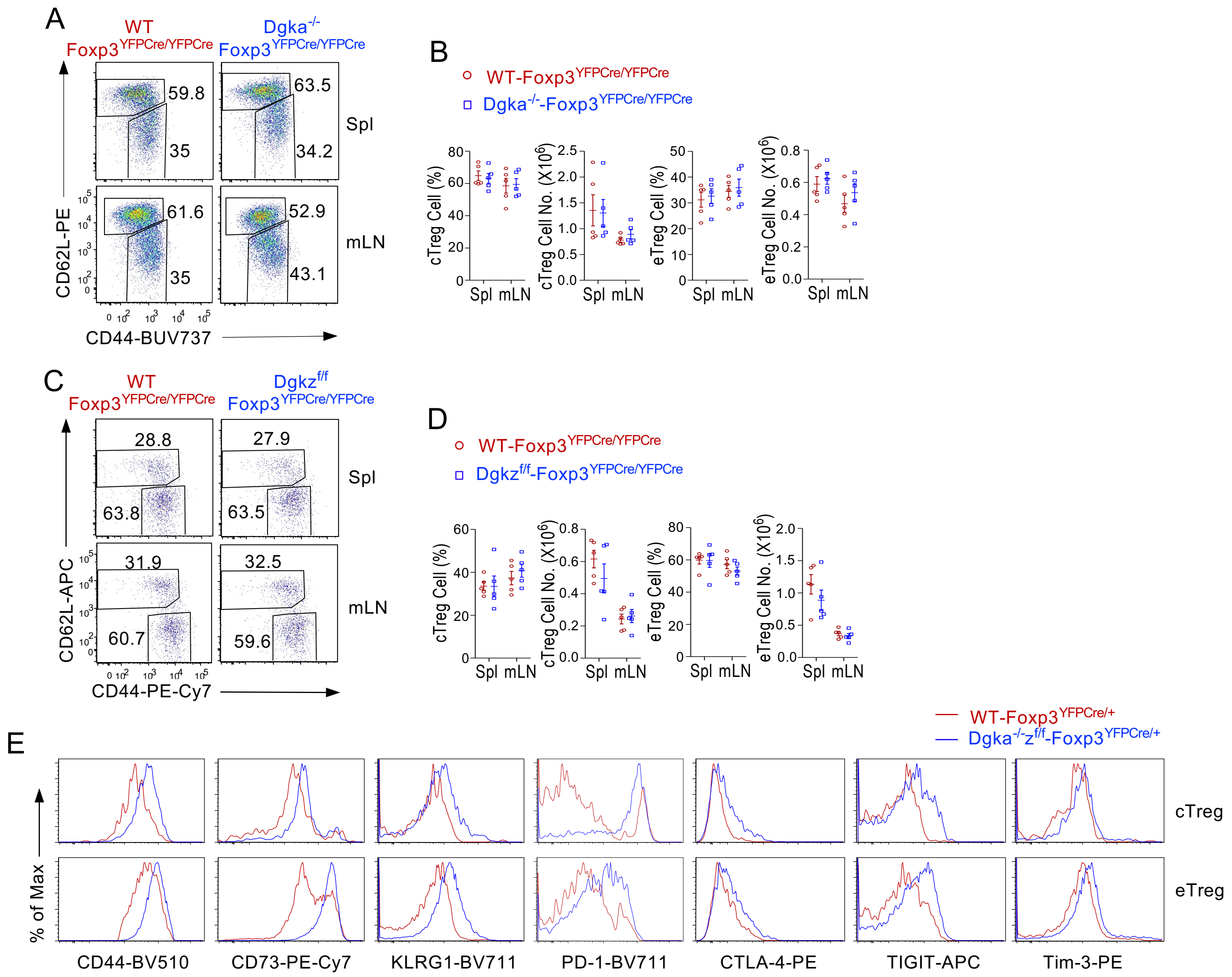


**Supplemental Figure S6. Treg properties in the absence of DGKα, ζ, or both. A, B.** Analyses of *Dgka*$^{-/-}$*z*$^{+/+}$*-Foxp3*$^{YFPCre/YFPCre}$ and WT-*Foxp3*$^{YFPCre/YFPCre}$ control mice. **A.** CD44 and CD62L expression in CD4$^{+}$Foxp3$^{+}$ Tregs. **B.** Scatter plots show mean ± SEM of cTreg and eTreg percentages and numbers. **C, D.** Analyses of *Dgka*$^{+/+}$*z*$^{f/f}$*-Foxp3*$^{YFPCre/YFPCre}$ and WT-*Foxp3*$^{YFPCre/YFPCre}$ control mice. **C.** CD44 and CD62L expression in CD4$^{+}$Foxp3$^{+}$ Tregs. **D.** Scatter plots show mean ± SEM of cTreg and eTreg percentages and numbers. **E.** Overlaid histograms comparing expression of indicated molecules in YFP$^{+}$ Tregs from female *Dgka*$^{-/-}$*z*$^{f/f}$*-Foxp3*$^{YFPCre/+}$ and WT-*Foxp3*$^{YFPCre/+}$ control mice. Data shown are presentative of or pooled from at least five experiments.
