## Supplemental figure S7 for "DGKα and ζ Deficiency Causes Regulatory T-Cell Dysregulation, Destabilization, and Conversion to Pathogenic T-Follicular Helper Cells to Trigger IgG1-Predominant Autoimmunity"

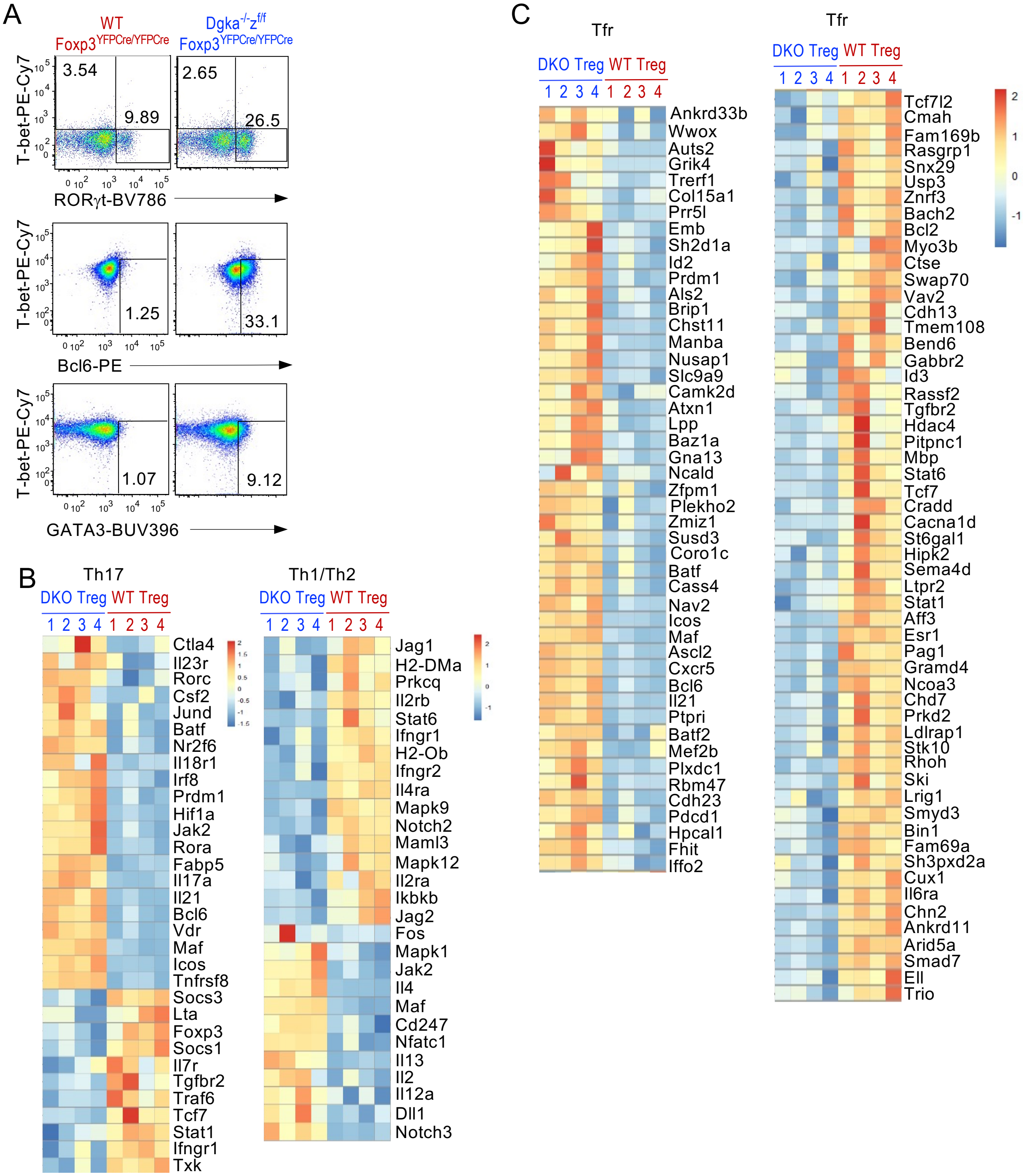


**Supplemental Figure S7. Analysis of effector lineages and gain-of-proinflammatory properties of αζDKO Tregs. A.** Intracellular staining of TF in mLN Tregs in WT-*Foxp3*$^{YFPCre/YFPCre}$ and *Dgka*$^{-/-}$*z*$^{f/f}$-*Foxp3*$^{YFPCre/YFPCre}$ mice. **B, C.** Heatmaps showing altered Th1, Th2, and Th17 cell (B) and Tfr cell (C) associated genes with statistically significant differences between WT and αζDKO Tregs ($p<0.05$).
