## Supplemental figure S8 for "DGKα and ζ Deficiency Causes Regulatory T-Cell Dysregulation, Destabilization, and Conversion to Pathogenic T-Follicular Helper Cells to Trigger IgG1-Predominant Autoimmunity"

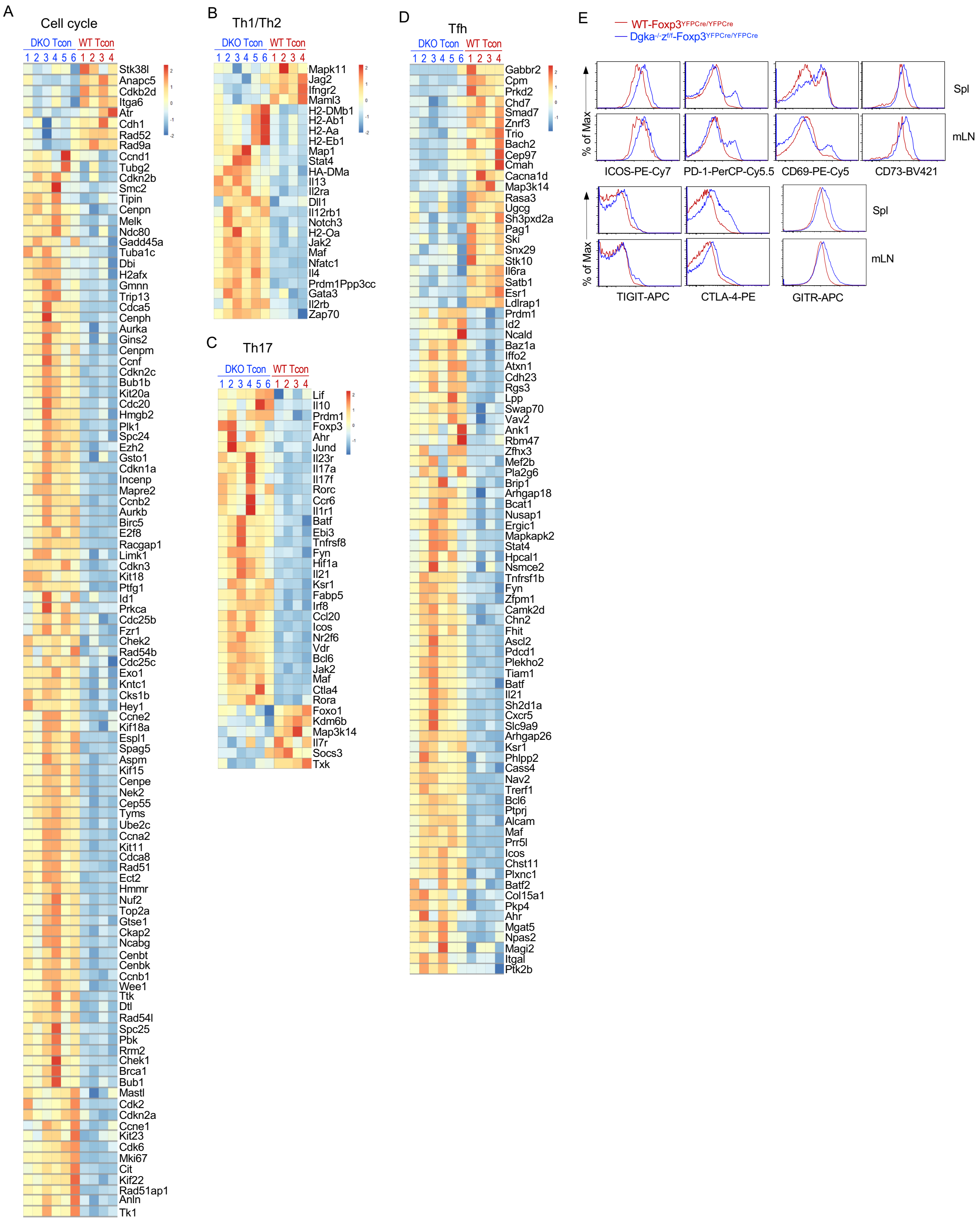


**Supplemental Figure S8. Altered cell cycle and Th associated pathways in CD4$^{+}$Foxp3$^{-}$ Tcons from Treg-αζDKO mice. A.** Heatmap shows DE cell cycle associated genes. **B.** Heatmap shows DE Th1/2 associated genes. **C.** Heatmap shows DE Th17 associated genes. **D.** Heatmap shows DE Tfh/Tfr associated genes. **E.** Alerted expression of surface molecules in Tcons from *Dgka$^{-/-}$z$^{f/f}$-Foxp3$^{YFPCre/YFPCre}$* and WT-*Foxp3$^{YFPCre/YFPCre}$* mice.
