## Supplemental figure S9 for "DGKα and ζ Deficiency Causes Regulatory T-Cell Dysregulation, Destabilization, and Conversion to Pathogenic T-Follicular Helper Cells to Trigger IgG1-Predominant Autoimmunity"

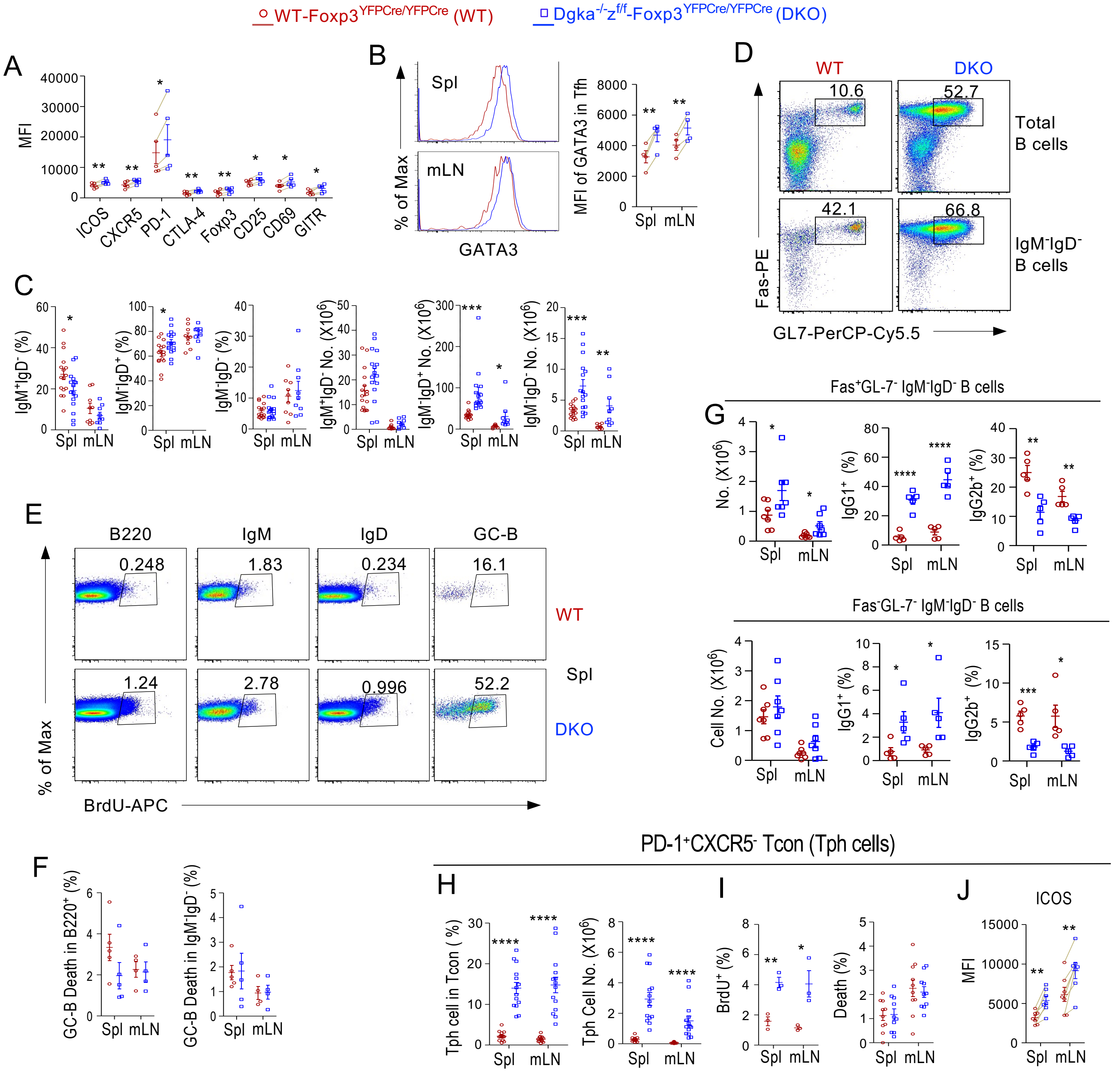


**Supplemental Figure S9. Assessment of Tfh, Tph and B cells in *Dgka$^{-/-}$z$^{f/f}$-Foxp3$^{YFPCre/YFPCre}$* mice. A.** Scatter plot shows mean ± SEM of MFI of the indicated molecules in Tfh cells. **B.** GATA3 levels in Tfh cells. **C.** IgM$^{+}$IgD$^{-}$, IgM$^{-}$IgD$^{+}$, and IgM$^{-}$IgD$^{-}$ B cells percentages and numbers. **D.** GC-B cell staining in mLNs. **E.** BrdU incorporation in B cell populations. **F.** Death rates of GC-B cells. **G.** Fas$^{+}$GL7$^{-}$ and Fas$^{-}$GL7$^{-}$ IgM$^{-}$IgD$^{-}$ B cell numbers and IgG1$^{+}$ or IgG2b$^{+}$ ratios. **H – J.** CD4$^{+}$TCRβ$^{+}$Foxp3$^{-}$PD1$^{+}$CXCR5$^{-}$ Tph cell percentages and numbers (H), BrdU incorporation and death rate (I), and ICOS levels (J). Each circle or square represents one mouse of the indicated genotypes. Data shown are presentative of or pooled from at least four experiments. *, p<0.05; **, p<0.01; ***, p<0.001; ****, p<0.0001 determined by two-tail unpaired Student t test (C, F, G – I) or pairwise Student t-test for A, B and J.
