## Supplemental figure S10 for "DGKα and ζ Deficiency Causes Regulatory T-Cell Dysregulation, Destabilization, and Conversion to Pathogenic T-Follicular Helper Cells to Trigger IgG1-Predominant Autoimmunity"

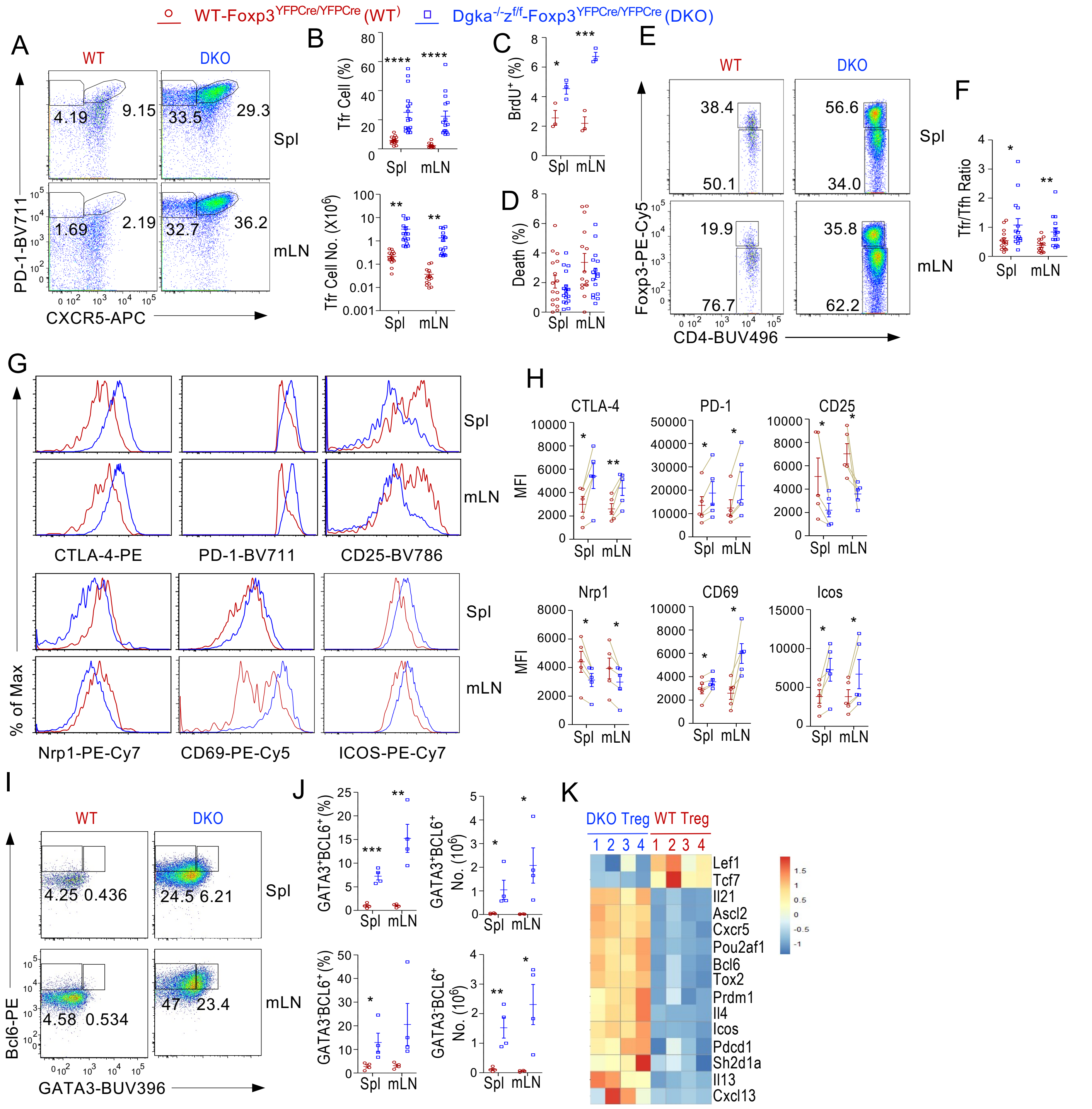


**Supplemental Figure S10. Dysregulation of Tfr cells in *Dgka$^{-/-}$z$^{f/f}$-Foxp3$^{YFPCre/YFPCre}$* mice.** Splenocytes and LN cells from *Dgka$^{-/-}$z$^{f/f}$-Foxp3$^{YFPCre/YFPCre}$* and WT-*Foxp3$^{YFPCre/YFPCre}$* mice were analyzed. **A.** CXCR5 and PD-1 expression in CD4$^{+}$Foxp3$^{+}$ Tregs. **B.** Mean ± SEM of Tfr percentages and numbers. **C.** BrdU incorporation in Tfr cells. **D.** Death rates of Tfr cells. **E.** Foxp3 and CD4 expression in CXCR5$^{+}$PD-1$^{+}$ CD4$^{+}$ T cells. **F.** Mean ± SEM of Tfr/Tcon ratios. **G.** Representative histograms show expression of indicated molecules in Tfr cells. **H.** Scatter plots represent mean ± SEM of MFI of the indicated molecules in Tfr cells. **I.** Bcl6 and GATA3 expression in CD4$^{+}$Foxp3$^{+}$ Tregs. **J.** Mean ± SEM of percentages and numbers of GATA3$^{+}$Bcl6$^{+}$ and GATA3$^{-}$Bcl6$^{+}$ cells in Tregs. **K.** Heatmap showing expression of Tfr-/Tfh-cell-related genes in WT and DKO Tregs. Data shown are representative of or pooled from 3–16 experiments. *, $p < 0.05$; **, $p < 0.01$; ***, $p < 0.001$; ****, $p < 0.0001$ determined by two-tail unpaired Student *t* test (B – D, F and J) or pairwise Student t-test for H.
