## Supplemental figure S11 for "DGKα and ζ Deficiency Causes Regulatory T-Cell Dysregulation, Destabilization, and Conversion to Pathogenic T-Follicular Helper Cells to Trigger IgG1-Predominant Autoimmunity"

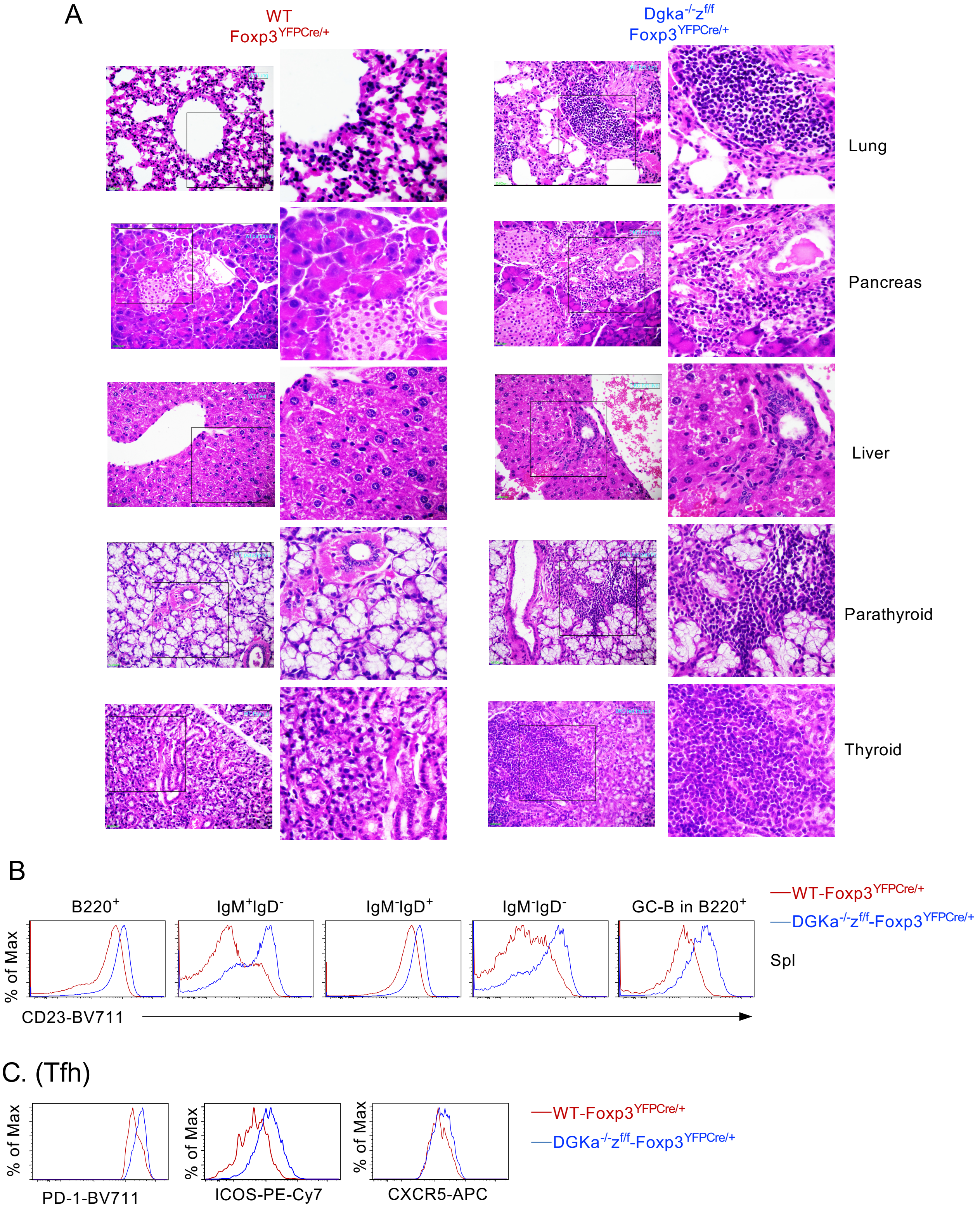


**Supplemental Figure S11. Development of autoimmune diseases in female *Dgka$^{-/-}$z$^{f/f}$-Foxp3$^{YFPCre/+}$* mice.** Three – nine months old female *Dgka$^{-/-}$z$^{f/f}$-Foxp3$^{YFPCre/+}$* (DKO-Cre$^{het}$) and WT-*Foxp3$^{YFPCre/+}$* (WT-Cre$^{het}$) mice were analyzed. **A.** Representative H&E staining of thin sections of the indicated organs. **B.** CD23 levels in the indicated B cell populations. **C.** PD-1, ICOS, and CXCR5 levels in Tfh cells.
