## Supplemental figure S12 for "DGKα and ζ Deficiency Causes Regulatory T-Cell Dysregulation, Destabilization, and Conversion to Pathogenic T-Follicular Helper Cells to Trigger IgG1-Predominant Autoimmunity"

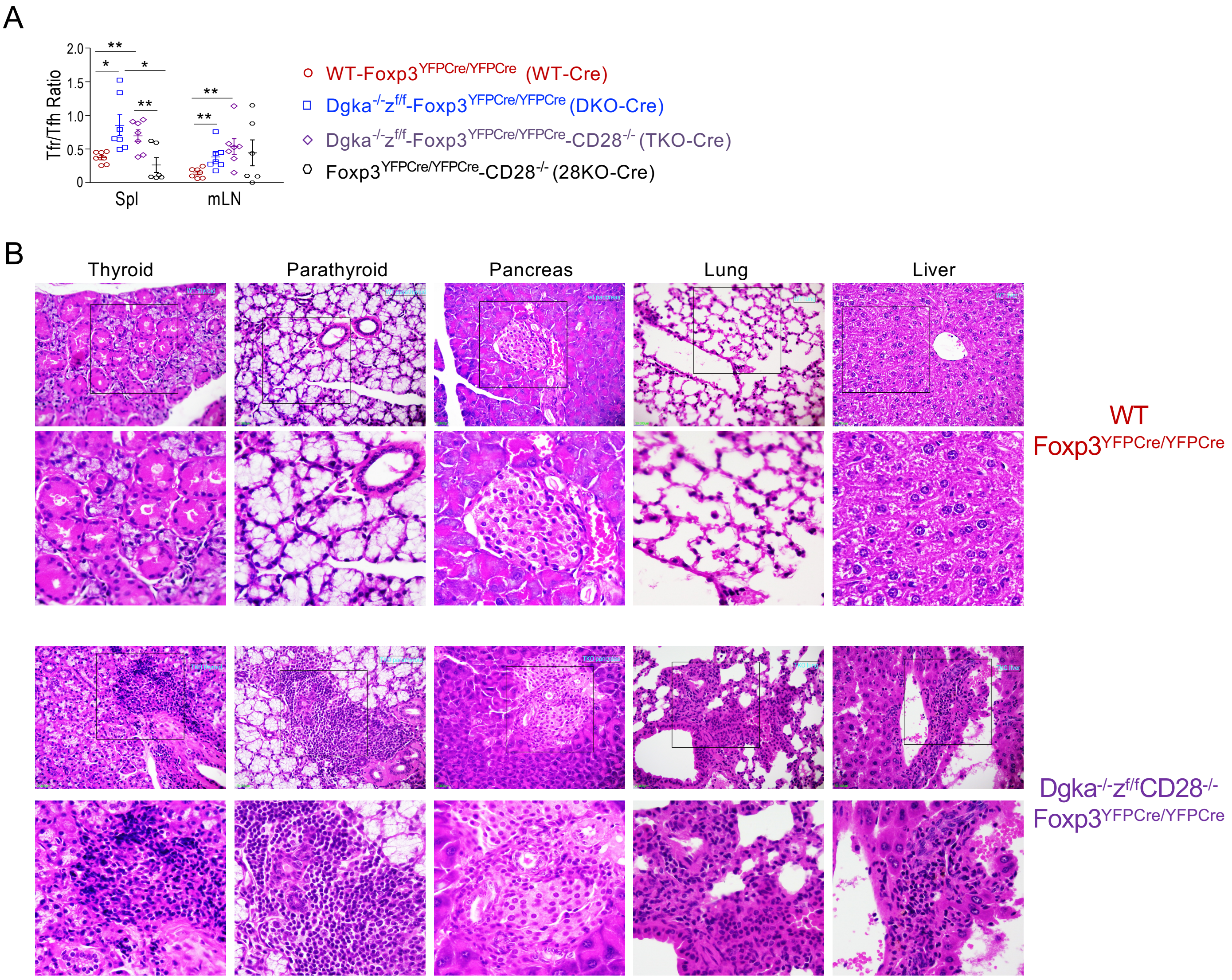


**Supplemental Figure S12. Altered Tfr/Tfh cell ratios and development of autoimmune diseases in *Dgka$^{-/-}$z$^{f/f}$-Foxp3$^{YFPCre/YFPCre}$-CD28$^{-/-}$* mice. A.** Tfr/Tfh cell ratios in the indicated mice. **B.** Representative H&E staining of tissue thin sections in the indicated mice are shown. Data shown are pooled from 5–10 experiments. *, $p < 0.05$; **, $p < 0.01$ determined by two-tail unpaired Student *t* test.
