## Supplemental figure S13 for "DGKα and ζ Deficiency Causes Regulatory T-Cell Dysregulation, Destabilization, and Conversion to Pathogenic T-Follicular Helper Cells to Trigger IgG1-Predominant Autoimmunity"

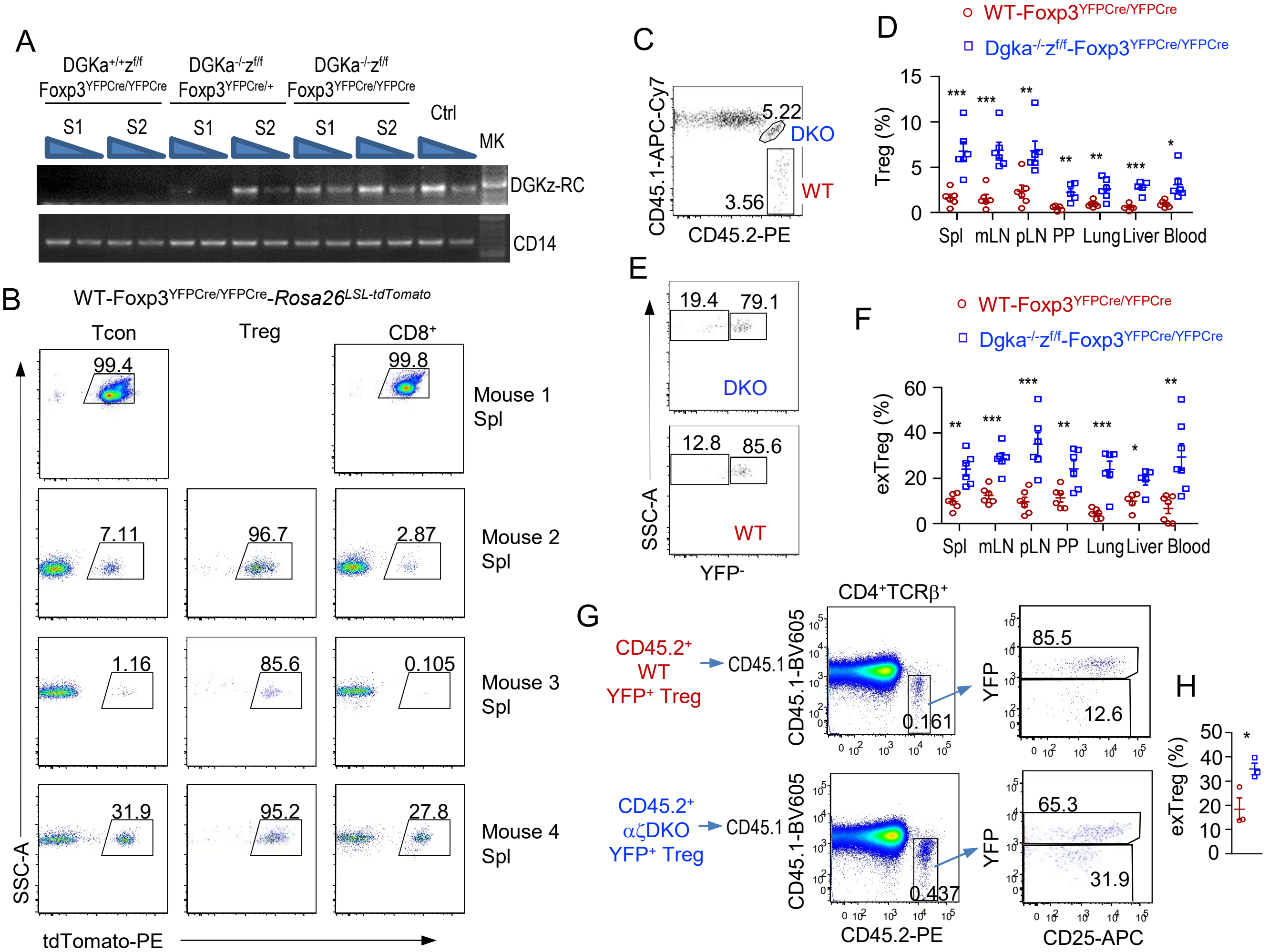


**Supplemental Figure S13. Enhanced αζDKO Treg conversion to exTregs/exTreg-Tfh cells. A.** Detection of Cre mediated recombination in CD4$^+$Foxp3YFP$^-$ Tcons in *Dgka*$^{-/-}$*z*$^{f/f}$*-Foxp3*$^{YFPCre/YFPCre}$ and *Dgka*$^{-/-}$*z*$^{f/f}$*-Foxp3*$^{YFPCre/+}$ mice but not in *Dgka*$^{+/+}$*z*$^{f/f}$*-Foxp3*$^{YFPCre/YFPCre}$ mice. Ctrl: DNA from total LN cells (including both Tregs and Tcons) of *Dgka*$^{-/-}$*z*$^{f/f}$*-Foxp3*$^{YFPCre/YFPCre}$ mice. MK: DNA ladder. S1 and S2 indicate different mice. One of two experiments of total four *Dgka*$^{+/+}$*z*$^{f/f}$*-Foxp3*$^{YFPCre/YFPCre}$ mice and four αζDKO mice is shown. The Cre mediated recombination of floxed Dgkz allele was detected with two Dgkz primers (forward: 5′-GGCACACTGGAGACTCTCAC-3′ and reverse: 5′-CTAGCTTGGGCTGCAGGT-3′). The CD14 genomic DNA was amplified using the following primers: Cd14forward: 5′-GCTCAAACTTTCAGAATCTACCGAC-3′ and Cd14reverse: 5′-AGTCAGTTCGTGGAGGCCGGAAATC-3′. **B.** Variegated tdTomato expression in *Foxp3*$^{YFPCre/YFPCre}$*-Rosa26*$^{LSL\text{-}tdTomato}$ mice**. C–F.** TCRβ$^{-/-}$ mice were coinjected with a mixture of CD45.1$^+$CD45.2$^-$ WT CD4$^+$ T cells, CD45.1$^-$CD45.2$^+$ CD4$^+$YFP$^+$ WT Tregs, and CD45.1$^+$CD45.2$^+$ CD4$^+$YFP$^+$ Treg-αζDKO Tregs and analyzed 8 weeks later. **C.** Representative FACS plots showing CD45.1 and CD45.2 staining in live gated mLN CD4$^+$TCRβ$^+$ cells. **D.** Percentages of WT and αζDKO Treg-derived cells within adoptively transferred CD4$^+$TCRβ$^+$ cells. **E.** FACS plots showing YFP levels in live gated cells derived from WT or αζDKO Tregs. **F.** Percentages of YFP$^-$ exTregs. Data shown are representative or pooled from 5–7 experiments. *, p<0.05; **, p<0.01; ***, p<0.001 determined by two-tail unpaired Student t-test. **G, H.** 5X10$^5$ Foxp3YFP$^+$ Tregs from WT-*Foxp3*$^{YFPCre/YFPCre}$ and *Dgka*$^{-/-}$*z*$^{f/f}$*-Foxp3*$^{YFPCre/YFPCre}$ mice were separately *i.v.* injected into CD45.1$^+$ C57BL6/J mice 4 hours after irradiation (400Rad). LNs from recipient mice were examined 14 days later. **G.** Representative FACS plots showing CD4$^+$TCRβ$^+$ T cells. **H.** Scatter plots show mean ± SEM of data pooled from three experiments. *, p<0.05 determined by two-tail unpaired Student t-test.
